# AIE nanoparticles camouflaged with tumor cell-derived exosomes for NIR-II imaging-guided photothermal therapy

**DOI:** 10.1101/2021.04.19.440457

**Authors:** Yirun Li, Xiaoxiao Fan, Yuanyuan Li, Runze Chen, Huwei Ni, Yiyin Zhang, Qiming Xia, Zhe Feng, Ben Zhong Tang, Jun Qian, Hui Lin

## Abstract

Nanoparticles (NPs) assisted photothermal therapy (PTT) is a promising cancer treatment modality and has attracted the attention of the scientific mainstream. However, developing NPs that exhibit efficient optical properties and specific tumor targeting capability simultaneously is difficult. Herein, we develop hybrid nanovesicles consisting of tumor cell-derived exosomes (EXO) and organic aggregation-induced emission (AIE) nanoparticles (TT3-*o*CB NP@EXOs) with enhanced second near-infrared (NIR-II, 900-1700 nm) fluorescence property and PTT functionality. Compared with TT3-*o*CB NPs, TT3-*o*CB NP@EXOs exhibit excellent biocompatibility, specific targeting ability *in vitro*, homing to homologous tumors *in vivo*, and prolonged circulation time. Furthermore, TT3-*o*CB NP@EXOs were utilized as biomimetic NPs for NIR-II fluorescence imaging-guided PTT of tumors, due to their high and stable photothermal conversion capacity under 808 nm irradiation. Therefore, the tumor cell-derived EXO/AIE NP hybrid nanovesicles may provide an alternative artificial targeting strategy for improving tumor diagnosis and PTT.

## 1. Introduction

Cancer represents one of the most dangerous threats to human health and has substantially high morbidity and mortality rates annually^[1]^. Traditional anticancer therapies include surgery, chemotherapy and radiotherapy and so on ^[2]^. Despite the continuous advances of anticancer drugs and techniques, tumor metastasis and recurrence remain the main causes of cancer-associated mortality^[3]^. Therefore, developing novel diagnostic and treatment approaches to enhance current therapeutic modalities are urgently needed.

Near-infrared (NIR) fluorescence imaging and NIR light-trigged therapy have gained increasing attention due to their high spatiotemporal resolution, low radiation and non-invasion ability^[4]^. With the development of probes and optical technologies, fluorescence imaging in the second near-infrared spectral region (NIR-II, 900-1700 nm) has displayed a better penetration depth and higher spatial resolution than that in the first near-infrared spectral region (NIR-I, 700-900 nm)^[5]^. NIR-II fluorescence imaging guided-photothermal therapy (PTT), a precise, noninvasive and localized therapeutic approach induced by NIR-II fluorescent probes, has experienced dramatic development^[6]^. These probes are capable of emitting fluorescence signals in NIR-II window and converting absorbed NIR irradiation energy into heat and increasing the local temperature to kill tumor cells^[7]^. To date, an increasing number of NIR-II fluorescent probes, including quantum dots^[8]^, carbon nanotubes^[9]^, and organic dyes^[10]^, have been applied in tumor treatments. Among them, organic fluorophores with aggregation-induced emission (AIE) characteristics have been developed as one of ideal materials due to their excellent photostability, biocompatibility and potentially low biotoxicity^[11]^. Thus, with the assistant of controllable NIR laser, NIR-II fluorescence imaging guided-PTT has shown great potential for clinical diagnosis and precise treatment, which can prevent normal tissue damage during therapy.

AIE NP-based tumor-targeting delivery systems have shown promising therapeutic efficacy in cancer via the enhanced permeability and retention (EPR) effect^[10b]^. However, the EPR effect of some AIE NPs is not sufficient. To increase their accumulation capacity in tumors, AIE NPs are usually functionalized with targeted antibodies, peptides or other biomolecules^[12]^. Nevertheless, various obstacles remain, such as their enhanced immune elimination and instability in blood vessels^[13]^. In recent years, biomimetic NPs have emerged as a promising approach to improve delivery efficiency by combining the functionalities of natural biomaterials with the engineering versatility of nanomaterials, such as red blood cell (RBC) membrane-camouflaged NPs^[14]^, natural killer (NK) cell membrane-camouflaged NPs^[15]^, tumor cell membrane-camouflaged NPs and hybrid membrane-camouflaged NPs^[16]^. Exosomes (EXO) are small vesicles (50-200 nm) released from multivesicular bodies following fusion with the plasma membrane and are produced by many different cells^[17]^. Tumor cell-derived EXO have many properties which make them promising candidates for targeting tumor cells, including a lack of immunogenicity, excellent biocompatibility, prolonged circulation owing to their endogenous origin, and a self-targeting homing ability to homologous tumors based on membrane-specific protein profiles^[18]^. Therefore, EXO can potentially enhance the tumor accumulation efficacy of AIE NPs, since the abundant proteins on the membrane surface can be inherited by EXO-camouflaged NPs^[19]^.

In our previous study, a novel organic fluorophore, TT3-*o*CB, was synthesized and encapsulated into nanoparticles (TT3-*o*CB NPs) to achieve high-resolution and real-time NIR-IIb fluorescence imaging of vessels and organs^[20]^. Recently, we found good photothermal property of this kind of AIE NPs *in vitro*. However, the poor tumor targeting capacity has limited further therapeutics development of the AIE NPs. To overcome this problem, we developed hybrid nanovesicles consisting of tumor cell-derived EXO and organic AIE NPs (TT3-*o*CB NP@EXOs) via electroporation. Compared with TT3-*o*CB NPs, TT3-*o*CB NP@EXOs exhibited greatly enhanced blood retention *in vivo*, as well as increased tumor uptake *in vitro* and *in vivo*. Because of their facile preparation process, bright NIR-II fluorescence emission, and high photothermal conversion capacity, TT3-*o*CB NP@EXOs could be utilized as biomimetic NPs for diagnostics and PTT. This combinatory strategy is an effective approach to the diagnosis and specifically targeted treatment of cancer, and has great potentials for the future clinical application of AIE luminogens.

## 2. Results and discussion

### 2.1. Preparation and characterization of TT3-*o*CB NPs@EXOs

TT3-*o*CB was prepared as a novel organic AIE fluorophore (Figure S1, Supporting Information), and its excellent AIE properties were thoroughly discussed in our previous study^[20]^. Briefly, TT3-*o*CB exhibited bright NIR-II fluorescence signal and good photothermal effect due to the molecular structure consisting of two components: first, a twisted architecture inherited from AIE that assured a high photoluminescent quantum yield (Φ_PL_) in aggregation state; second, an enhanced π-conjugated planar unit that could provide high molar extinction coefficient (ε)^[20]^. In this study, we encapsulated TT3-*o*CB into NPs by using the FDA-approved amphiphilic polymer, Pluronic F-127 (F-127), to improve its solubility in aqueous solution. EXO mediate intercellular communication and transfer various types of cargo. They have been proven to be a better natural delivery nanoplatform than traditional lipid transport proteins or lipoproteins^[21]^. Tumor cell-derived EXO were isolated from the supernatant of CT26 cells (murine colorectal cancer cell line) via differential ultracentrifugation^[22]^, and then electroporated with the existence of TT3-*o*CB NPs to obtain TT3-*o*CB NP@EXO hybrid nanovesicles (**Figure 1**). To explore whether the TT3-*o*CB NPs were sheathed with EXO, transmission electron microscopy (TEM), dynamic light scattering (DLS), zeta potential and western blot analyses were performed. The purified EXO had a cup-like shape, while both TT3-*o*CB NPs and TT3-*o*CB NP@EXOs had typical round shapes under TEM (**Figure 2A**). Their mean diameters were approximately 116 nm, 89 nm and 164 nm, respectively (Figure 2B). The zeta potential values of EXO, TT3-*o*CB NPs and TT3-*o*CB NP@EXOs were −8.5 ± 0.8 mV, −0.6 ± 1.1 mV and −6.4 ± 0.9 mV, respectively (Figure 2C). Western blot experiments further showed that the EXO markers TSG101, CD9 and CD81 were detectable in the purified EXO and TT3-*o*CB NP@EXOs rather than in the whole-cell lysates (Figure 2D), confirming the presence of EXO in TT3-*o*CB NP@EXOs. To investigate whether the EXO would affect the optical properties of TT3-*o*CB NPs, we examined the absorption and photoluminescence (PL) spectra. The absorption peaks of TT3-*o*CB NP@EXO hybrid nanovesicles were similar to those of TT3-*o*CB NPs, occurring at approximately 310 nm, 390 nm and 775 nm (same concentration: 0.1 mg mL^-1^, Figure 2E). Regarding their fluorescence properties, both curves of the PL spectra were in good agreement (same concentration: 1 mg mL^-1^, Figure 2F). Together, the above results support that we have successfully prepared the TT3-*o*CB NP@EXO hybrid nanovesicles and found that the good optical properties of TT3-*o*CB NPs would not be affected by the tumor cell derived-EXO.

**Figure 1.**
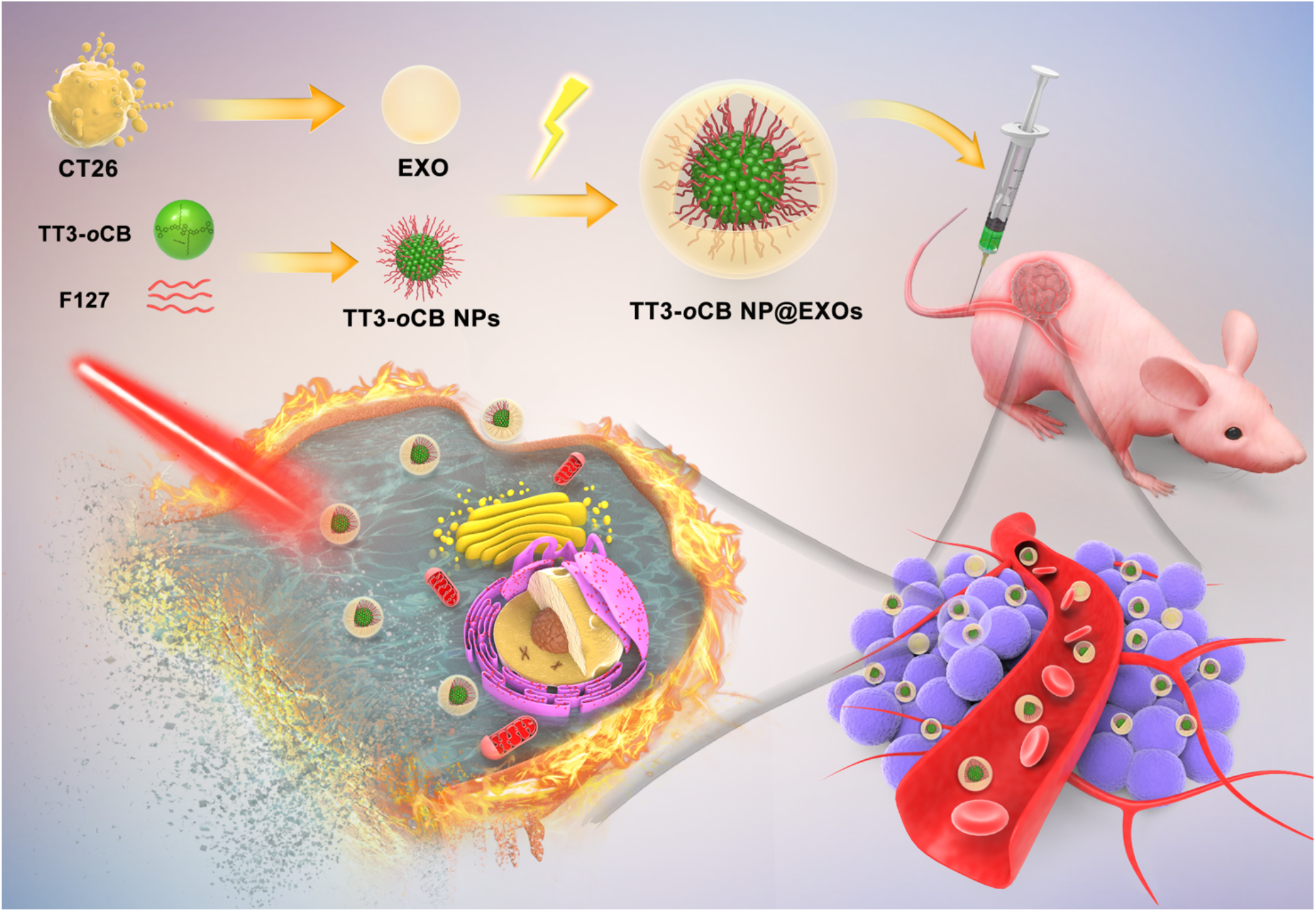
Schematic illustration of tumor cell-derived EXO/organic AIE NP hybrid nanovesicles which exhibit tumor-homing penetration, enhanced NIR-II fluorescence signals and efficient photothermal therapy abilities.

**Figure 2.**
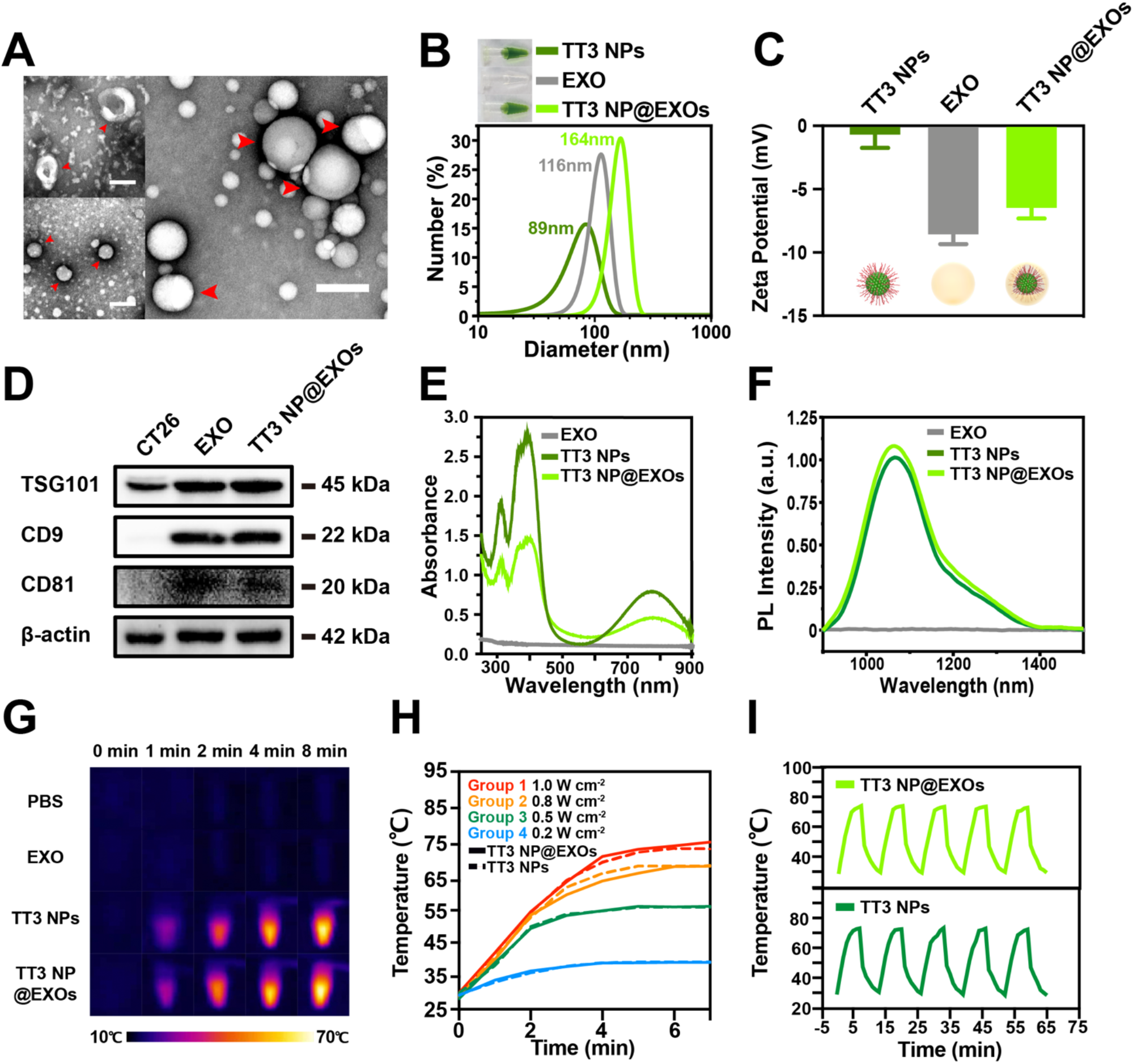
Physiochemical features of EXO, TT3-*o*CB NPs and TT3-*o*CB NP@EXOs in aqueous dispersions (termed TT3 NPs and TT3 NP@EXOs in figure, respectively). (A) TEM images of EXO, TT3-*o*CB NPs and TT3-*o*CB NP@EXOs (upper left panel: the red arrows indicate EXO isolated from CT26 cells; bottom left panel: the red arrows indicate TT3-*o*CB NPs; right panel: the red arrows indicate TT3-*o*CB NP@EXOs). DLS (B) and zeta potential (C) analyses of EXOs, TT3-*o*CB NPs and TT3-*o*CB NP@EXOs in dispersion (0.1 mg mL^-1^). (D) Western blot analyses of the EXO markers TSG101, CD9 and CD81 in the colorectal cancer cell line CT26, EXO isolated from CT26 cells and TT3-*o*CB NP@EXO hybrid nanovesicles. Absorption (E) and fluorescence emission spectra (F) of EXO, TT3-*o*CB NPs and TT3-*o*CB NP@EXOs in aqueous dispersion at the same concentration (EXO was measured by BCA assays). (G) Infrared thermal images of PBS, EXO, TT3-*o*CB NPs and TT3-*o*CB NP@EXOs in aqueous dispersion (0.1 mg mL^-1^) under irradiation of an 808 nm laser (1 W cm^-2^) for 1 min, 2 min, 4 min and 8 min. (H) Temperature curves of TT3-*o*CB NPs and TT3-*o*CB NP@EXOs in aqueous dispersion (0.1 mg mL^-1^) under irradiation of the 808 nm laser at different power densities (dashed lines: TT3-*o*CB NP solutions, solid lines: TT3-*o*CB NP@EXO solutions). (I) Photothermal stability of TT3-*o*CB NPs and TT3-*o*CB NP@EXOs (808 nm laser, 1 W cm^-2^, 5 cycles).

To further examine the photothermal properties of TT3-*o*CB NPs and TT3-*o*CB NP@EXOs, the temperature of heating aqueous dispersions under continuous laser irradiation (808 nm, 1.0 W cm^-2^) was monitored at 0 min, 1 min, 2 min, 4 min and 8 min using an infrared thermal imaging camera. The temperature of the TT3-*o*CB NP@EXO dispersion (0.5 mg mL^-1^) and TT3-*o*CB NP dispersion (0.5 mg mL^-1^) increased by approximately 70 °C after irradiation for 8 min, but the control solutions (PBS and EXO) were hardly heated by irradiation (Figure 2G). Then, TT3-*o*CB NP and TT3-*o*CB NP@EXO aqueous dispersions were exposed to irradiation with the 808 nm laser at different power densities to investigate the photothermal performances. The temperatures of both TT3-*o*CB NP@EXOs and TT3-*o*CB NPs rapidly increased and gradually reached a plateau after approximately 4 min of laser irradiation. As shown in Figure 2H, the temperature increased to ∼73 °C when the power density was increased to 1.0 W cm^-2^, indicating that heat generation was regulated precisely and that the EXO membrane did not affect the photothermal efficiency. The photothermal conversion efficiency (*η*) of AIE NPs has been recognized as another significant factor underlying photothermal efficiency in cancer treatment. Photothermal conversion efficiencies of TT3-*o*CB NPs and TT3-*o*CB NP@EXOs were 34.0% and 34.7%, respectively, as determined by a previously described method^[23]^, and were comparable with some reported materials^[10b, 24]^. In addition, both aqueous dispersions (0.5 mg mL^-1^) were exposed to cyclic irradiation to evaluate their photothermal stability (808 nm, 1.0 W cm^-2^, 5 cycles). There was almost no change in their photothermal performances after five heating-cooling cycles, showing the high photothermal stability of both materials (Figure 2I). Overall, the TT3-*o*CB NP@EXO hybrid nanovesicles exhibited excellent photothermal performances, which were not affected by EXO.

### 2.2. Verify specific tumor targeting of TT3-*o*CB NP@EXOs using NIR-II fluorescence microscopy

Previous studies found that EXO exhibited the capabilities of interacting with and targeting homologous cells dependent on the surface adhesion proteins and vector ligands, which could be used as specific delivery vehicles for various therapeutic agents^[25]^. Based on these capabilities of EXO, we performed colocalization imaging to explore whether the CT26 cell derived-EXO camouflaged-TT3-*o*CB NPs could exhibit enhanced targeting ability in CT26 cells rather than in Hep1-6 (murine liver cancer cell line) or NIH3T3 cells (murine fibroblast cell line). As shown in **Figure 3**, after incubation with EXO, TT3-*o*CB NPs and TT3-*o*CB NP@EXOs for 24 hours, the three cell lines exhibited obviously different results. Upon 808 nm laser irradiation, strong NIR-II fluorescence signals consistent with distinct accumulation of TT3-*o*CB NP@EXOs were observed in CT26 cells, while only a small amount of NIR-II fluorescence signals were observed in CT26 cells treated with TT3-*o*CB NPs (Figure 3A). As shown in Figure 3B and 3C, there was no difference in NIR-II fluorescence intensities between the TT3-*o*CB NPs and TT3-*o*CB NP@EXOs in either Hep1-6 or NIH3T3 cells. However, NPs are usually captured and accumulated in liver^[26]^. The liver-originated cancer cell line Hep1-6 could capture more NPs than the fibroblast cell line NIH3T3. These results confirmed that EXO was a promising platform to deliver NPs to homologous tumor cells, and the EXO camouflaged-TT3-*o*CB NPs can be used for NIR-II fluorescence tumor imaging due to their tumor cell-specific penetration ability.

**Figure 3.**
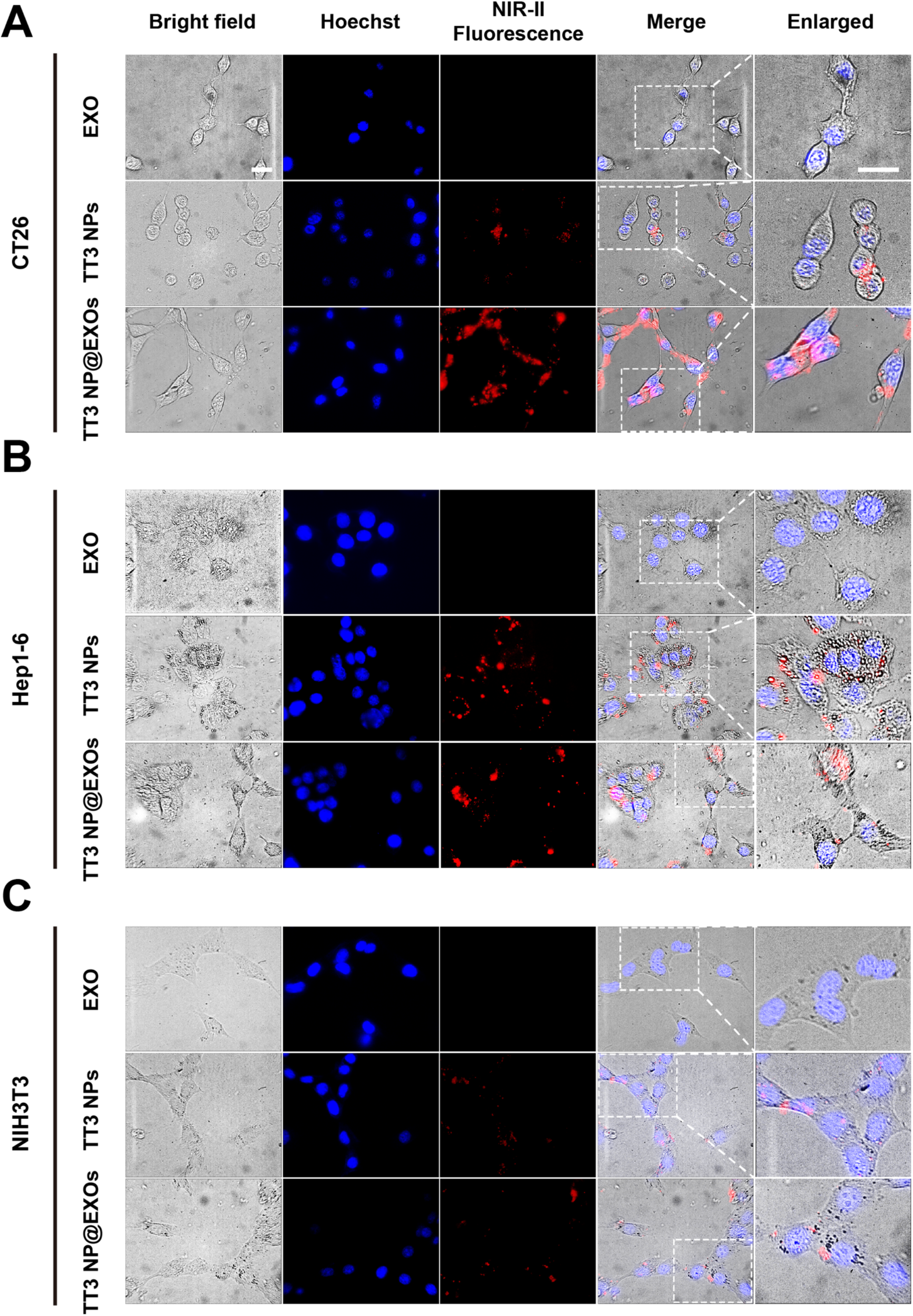
Bright field and fluorescence microscopic images of different cell lines. NIR-II fluorescence microscopic images of the colorectal cancer cell line CT26 (A), mouse liver cancer cell line Hep1-6 (B) and fibroblast cell line NIH3T3 (C) were acquired after incubation with EXO, TT3-*o*CB NPs and TT3-*o*CB NP@EXOs (50 mg mL^-1^, termed TT3 NPs and TT3 NP@EXOs), respectively. All the cell lines were stained with Hoechst, which emitted blue fluorescence signals. Red: NIR-II fluorescence from TT3-*o*CB NPs or TT3-*o*CB NP@EXOs), blue: Hoechst-stained cell nuclei. Scale bar: 20 μm.

### 2.3. *In vitro* cytotoxicity and tumor-targeting PTT assays

The *in vitro* toxicity of TT3-*o*CB NPs and TT3-*o*CB NP@EXOs to different cell lines (CT26, Hep1-6 and NIH3T3) was examined by MTS assays. Both materials showed negligible cytotoxicity on the three cell lines even at a high concentration of 50 μg mL^-1^, and the cytotoxicities of the materials were not significantly different (**Figure 4**A; Figure S2, Supporting Information). The proteins on the membranes of tumor cell-derived EXO generally govern their specific homing to and selective entry into homologous tumor cells^[18a]^. To confirm the ability of CT26 cell derived-EXO camouflaged-TT3-*o*CB NPs to be internalized specifically by CT26 cells and the photothermal efficiency *in vitro*, we selected three different cell lines, irradiated them by an 808 nm laser for 5 min (1.0 W cm^-2^) and then analyzed the viability by MTS assays. The PTT efficiency of TT3-*o*CB NP@EXOs in CT26 cells was markedly higher compared with that of TT3-*o*CB NPs (Figure 4B), which was consistent with the NIR-II fluorescence imaging data shown above (Figure 3A). For Hep1-6 and NIH3T3, there was no difference in cell viability between the TT3-*o*CB NPs and TT3-*o*CB NP@EXOs (Figure 4B). Then, colony formation and wound healing assays were performed to assess the phototoxic effects on cell proliferation and migration in the different groups. Similarly, colony formation assays revealed that the clonogenic survival of CT26 cells treated with TT3-*o*CB NP@EXOs and irradiation was significantly decreased compared with that of TT3-*o*CB NPs. No obvious differences in the clonogenic survival of Hep1-6 and NIH3T3 cells were observed between the TT3-*o*CB NP and TT3-*o*CB NP@EXO treated groups (Figure 4C). Next, wound healing assays showed that TT3-*o*CB NP@EXOs significantly inhibited the migration of CT26 cells compared to the PBS, EXO and TT3-*o*CB NPs, after 808 nm laser irradiation for 5 min (Figure S3A, Supporting Information). There was no difference in the migration of Hep1-6 cells between the TT3-*o*CB NP and TT3-*o*CB NP@EXO treated groups, while the wound healing areas were not different among all four treated NIH3T3 cell groups (Figure S3B and S3C, Supporting Information). *In vitro* experiments revealed that CT26-derived EXO enhanced the specific penetration and PTT efficiency of TT3-*o*CB NPs in CT26 cells but not in other tumor or normal cells.

**Figure 4.**
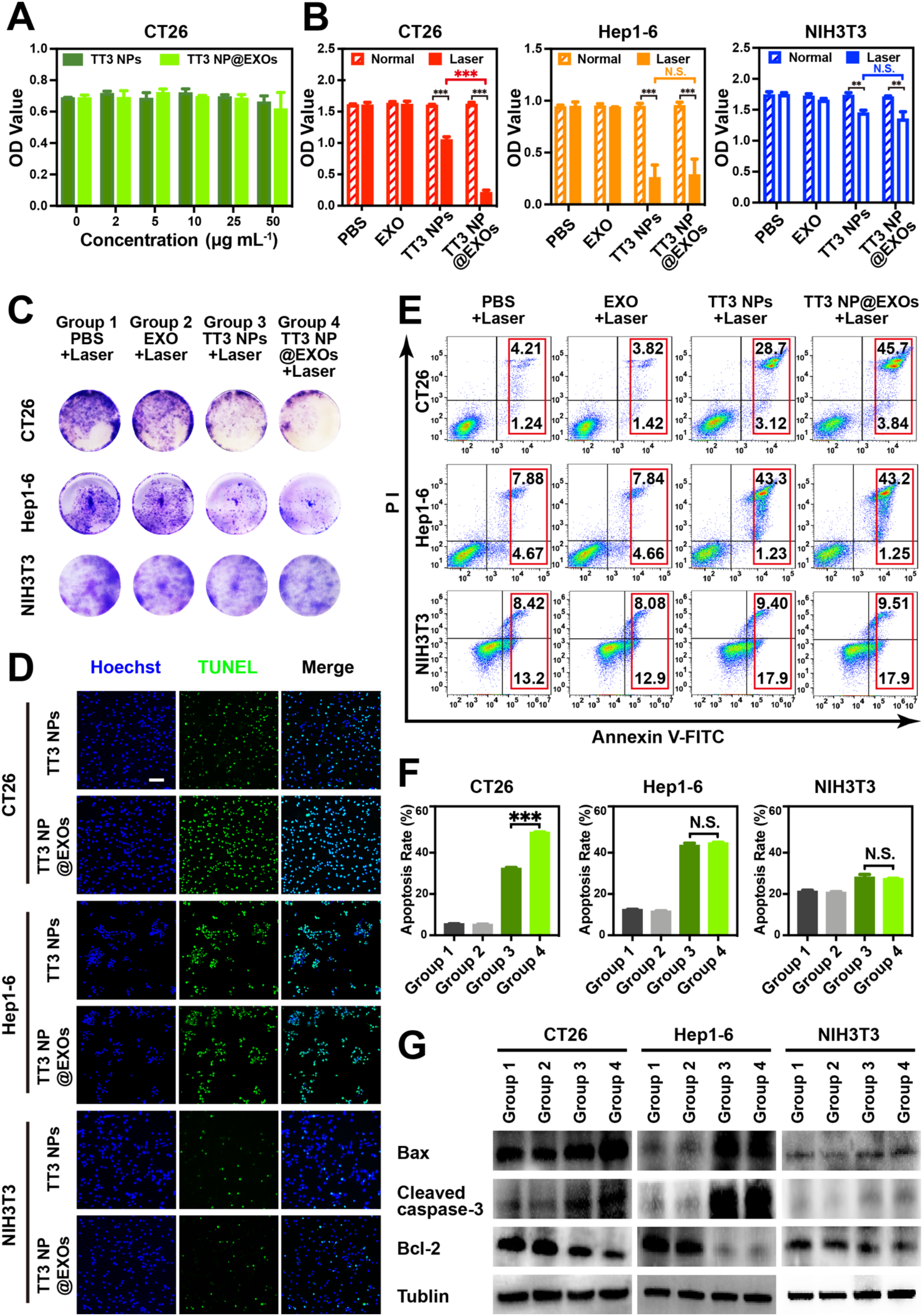
*In vitro* PTT effects of TT3-*o*CB NPs and TT3-*o*CB NP@EXOs on different cell lines (termed TT3 NPs and TT3 NP@EXOs, respectively). (A) *In vitro* viability of CT26 cells incubated with TT3-*o*CB NPs and TT3-*o*CB NP@EXOs at different concentrations. (B) Viabilities of CT26, Hep1-6 and NIH3T3 cells incubated with PBS, EXO, TT3-*o*CB NPs and TT3-*o*CB NP@EXOs, with and without the irradiation of 808 nm laser (1.0 W cm^-2^, 5 min, ** *P* < 0.01, *** *P* < 0.001). (C) Clone formation assays of CT26, Hep1-6 and NIH3T3 cells incubated with PBS, EXO, TT3-*o*CB NPs and TT3-*o*CB NP@EXOs under the irradiation of 808 nm laser (1.0 W cm^-2^, 5 min). (D) TUNEL staining of CT26, Hep1-6 and NIH3T3 cells incubated with TT3-*o*CB NPs and TT3-*o*CB NP@EXOs, under the irradiation of 808 nm laser (1.0 W cm^-2^, 5 min; green: TUNEL-positive cells, blue: Hoechst-stained cell nuclei). Scale bar: 50 μm. (E, F) Flow cytometry analyses of CT26, Hep1-6 and NIH3T3 cells incubated with PBS, EXO, TT3-*o*CB NPs and TT3-*o*CB NP@EXOs, under the irradiation of 808 nm laser (1.0 W cm^-2^, 5 min). Annexin V-FITC/PI staining was used (n=3, N.S. *P* > 0.05, *** *P* < 0.001). (G) Western blot analyses of apoptosis markers in CT26, Hep1-6 and NIH3T3 cells incubated with PBS (group 1), EXO (group 2), TT3-*o*CB NPs (group 3) and TT3-*o*CB NP@EXOs (group 4), under the irradiation of 808 nm laser (1 W cm^-2^, 5 min).

### 2.4. Anticancer activity through regulating cell apoptosis *in vitro*

As most of the PTT materials used for cancer therapy are believed to trigger apoptosis^[27]^, we hypothesized that the PTT of TT3-*o*CB NPs/TT3-*o*CB NP@EXOs would also initiate the apoptosis of tumor cells. After PTT, the apoptosis of different cell lines was detected by the terminal deoxynucleotidyl transferase-mediated dUTP nick-end labeling (TUNEL) assay. The proportion of representative apoptosis-positive CT26 cells in the TT3-*o*CB NP@EXO&808 nm laser treated group was obviously higher than those in the other three groups, while no differences in Hep1-6 and NIH3T3 cell apoptosis were observed between the TT3-*o*CB NP and TT3-*o*CB NP@EXO treated groups (Figure 4D; Figure S4, Supporting Information). TT3-*o*CB NPs&808 nm irradiation had killing effect on tumor cells Hep1-6 rather than normal cells NIH3T3, but it could not be improved by CT26 derived-EXO. Next, flow cytometry was used to determine the effects of different materials on cell apoptosis. As shown in Figure 4E and 4F, the percentage of total apoptotic CT26 cells in the TT3-*o*CB NP@EXO&808 nm laser treated group was enhanced compared to that in the TT3-*o*CB NP&808 nm laser treated group (49.82 ± 0.27% vs. 32.39 ± 0.50%). For Hep1-6 and NIH3T3 cells, no significant differences in the percentages of total apoptotic cells were observed between the TT3-*o*CB NP&irradiation and TT3-*o*CB NP@EXO&irradiation treated groups (Hep1-6: 43.46 ± 1.18% vs. 44.69 ± 0.37%; NIH3T3: 28.03 ± 1.45% vs. 27.35 ± 0.20%). To further explore the mechanism by which PTT induced the apoptosis of tumor cells, we evaluated the key apoptosis-related proteins Bax, cleaved caspase-3 and Bcl-2 by Western blot, and observed higher expression of Bax and cleaved caspase-3 and lower expression of Bcl-2 in TT3-*o*CB NP@EXO&irradiation treated CT26 cells, compared to TT3-*o*CB NP&irradiation treated CT26 cells. Consistent with the above results, no differences in the expression levels of the abovementioned apoptosis-related markers in Hep1-6 or NIH3T3 cells were observed between the two groups (Figure 4G). These results demonstrated that PTT induced the apoptosis of tumor cells, which could be a major mechanism by which TT3-*o*CB NPs and TT3-*o*CB NP@EXOs treat cancer.

### 2.5. Biodistribution of TT3-*o*CB NP@EXOs in CT26 tumor-bearing mice

After elucidating the anticancer effect of TT3-*o*CB NP@EXOs *in vitro*, we next addressed the *in vivo* behavior and fate of TT3-*o*CB NP@EXOs in a CT26 tumor-bearing mouse model with a lab-built NIR-II fluorescence *in vivo* imaging system. Upon the intravenous (i.v.) injection of 200 μL aqueous dispersion of TT3-*o*CB NPs or TT3-*o*CB NP@EXOs (1 mg mL^-1^) into mice, noninvasive NIR-II fluorescence imaging was performed to track the distributions of TT3-*o*CB NPs and TT3-*o*CB NP@EXOs 24 h after the injection. As shown in **Figure 5**A, significant fluorescence signals were observed in the tumors of the TT3-*o*CB NP@EXO treated group, while no obvious fluorescence signals were detected in the tumors of the TT3-*o*CB NP treated group. The above result demonstrated that EXO could confer AIE NPs, which cannot selectively target tumors, a specific tumor targeting capability. We speculated that there may be two reasons. First, compared to the NPs alone, more tumor cell derived-EXO-camouflaged NPs might be taken up by tumor cells due to the homing ability of EXO, which we have verified *in vitro*. Second, the EXO might improve the blood circulation time of NPs. Therefore, we extracted blood samples from mice at designated time intervals after i.v. injection and examined the NIR-II fluorescence signals. We found that although the NIR-II fluorescence signals decreased gradually in the blood samples of both groups, TT3-*o*CB NP@EXOs circulated for a longer period than the TT3-*o*CB NPs, as expected (Figure 5B). The EXO might prolong the NPs’ circulation time due to their delayed capture by the liver. For verification, we performed the time-dependent biodistribution assay to further investigated the distributions of materials *in vivo*. The tumors and vital organs were collected from sacrificed mice at designated time intervals (12 h, 24 h, 48 h and 72 h). As shown in Figure 5C and 5D, the fluorescence signals substantially accumulated in the liver and spleen for elimination by phagocytosis in the reticuloendothelial system^[18b]^. We further analyzed the intensity of fluorescence signals in livers between two groups. In general, the fluorescence signals in livers of TT3-*o*CB NP@EXO treated group were lower than those of the TT3-*o*CB NP treated group at all four time points post injection. Among them, the most significant difference was observed at 24 h post injection (*P* < 0.05), revealing that the EXO reduced the capture of NPs by the livers (Figure S5, Supporting Information). For other vital organs, all the fluorescence signals in the organs peaked at 24 h post injection and decreased over time (Figure 5C and 5D). Biodistribution in tumors was also assessed by *ex vivo* fluorescence imaging, revealing that the fluorescence signal in tumors of the TT3-*o*CB NP@EXO treated group peaked at 24 h post injection and decreased over time, while the fluorescence signal was barely detectable in the tumors of the TT3-*o*CB NP treated group, consistent with previous observation (Figure 5A, 5C, 5D; Figure S6, Supporting Information). Quantitative analyses of the tumors illustrated fluorescence signals in the TT3-*o*CB NP@EXO treated group were approximately 5.9- and 4.3-fold higher than those in the TT3-*o*CB NP treated group at 24 and 48 h post injection, respectively (Figure 5E).

**Figure 5.**
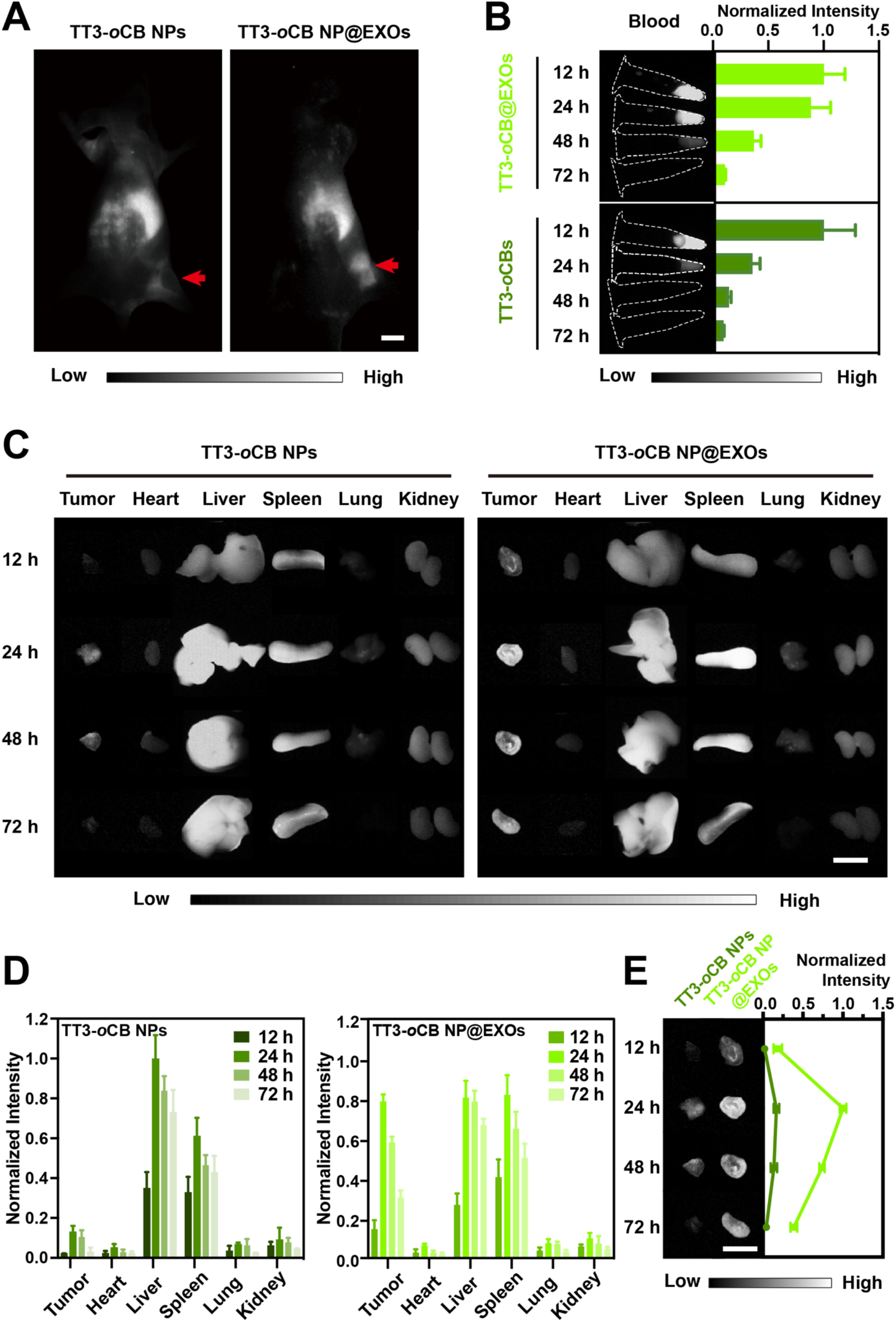
*In vivo* biodistribution of TT3-*o*CB NPs and TT3-*o*CB NP@EXOs. (A) Representative NIR-II fluorescence images of CT26 tumor-bearing mice 24 h after the intravenous (i.v.) administration of TT3-*o*CB NPs (left) and TT3-*o*CB NP@EXOs (right). Scale bar: 10 mm. Intensity of 808 nm laser excitation: 50 mW/cm^2^. (B) Time-dependent NIR-II fluorescence images (left) and normalized intensity changes (right) of blood extracted from TT3-*o*CB NP-treated and TT3-*o*CB NP@EXO-treated CT26 tumor-bearing mice, 12, 24, 48, and 72 h post injection (1 mg mL^-1^, 200 μL, n = 3, intensity of 808 nm laser excitation: 50 mW/cm^2^.). *Ex vivo* NIR-II fluorescence images (C) and normalized NIR-II fluorescence intensities (D) of the CT26 tumors and major organs of mice i.v. injected with TT3-*o*CB NPs (left) and TT3-*o*CB NP@EXOs (right), 12, 24, 48, and 72 h post injection (n = 3). Scale bar: 10 mm. Intensity of 808 nm laser excitation: 50 mW/cm^2^. (E) Time-dependent NIR-II fluorescence images (left) and normalized intensity changes (right) of the tumors in the TT3-*o*CB NP- and TT3-*o*CB NP@EXO-treated groups, 12, 24, 48, and 72 h post injection (n = 3). Scale bar: 10 mm. Intensity of 808 nm laser excitation: 50 mW/cm^2^.

Moreover, considering that the TT3-*o*CB NP@EXOs could successfully target to tumors for 24 h, we further verified their tracking capacity in CT26 tumor-bearing mice. Remarkably high fluorescence brightness was observed in the tumors of the TT3-*o*CB NP@EXO treated group on day 1 post injection, and the brightness decreased over time. Finally, no fluorescence signal was detectable in tumors on day 15 (Figure S7, Supporting Information), as the persistent differentiation and proliferation of tumor cells induced the metabolism and excretion of the NPs. The above biodistribution results obtained *in vivo* and *ex vivo* inspired us to further study the potential of TT3-*o*CB NP@EXOs for the precise targeting and PTT of tumors *in vivo*.

### 2.6. *In vivo* PTT

BALB/c nude mice (4 weeks old) subcutaneously injected with CT26 cells were used as the tumor model (**Figure 6**A). When the tumor volumes reached approximately 40–60 mm^3^, forty mice were randomly assigned to eight treatment groups: (I) PBS, (II) PBS+laser, (III) EXO, (IV) EXO+laser, (V) TT3-*o*CB NPs, (VI) TT3-*o*CB NPs+laser, (VII) TT3-*o*CB NP@EXOs, and (VIII) TT3-*o*CB NP@EXOs+laser (n = 5, i.v. concentration: 1 mg mL^-1^, 200 μL, 808 nm irradiation power: 1.0 W cm^-2^, 10 min, Figure 6A; Figure S8, Supporting Information). As shown in Figure 6B and 6C, the temperature in the tumor region of the TT3-*o*CB NP@EXO+laser-treated mice rapidly rose to ∼ 57 °C in 10 min (beyond the threshold tolerated by cancer cells), compared to ∼ 47 °C in mice treated with TT3-*o*CB NPs+laser, ∼ 41 °C in mice treated with EXO+laser and ∼ 41 °C in mice treated with PBS+laser. After irradiation, the animal body weights, tumor volumes and tumor weights were recorded every two days. There were no dramatic changes in body weight 9 days post irradiation (Figure 6D). After 15 days, all the mice were sacrificed, and the tumors were extracted and weighed. Only the tumors of the mice treated with TT3-*o*CB NP@EXOs+laser were obviously inhibited or even eradicated compared to the tumors in all the other groups (Figure 6E and 6F). The tumors of mice were harvested for photograph, as well as histological analyses by H&E and TUNEL assays (Figure 6H; Figure S9, Supporting Information). The most severe damage (nuclear fragmentation and apoptosis) was observed in the group treated with TT3-*o*CB NP@EXOs+laser. These findings confirmed that TT3-*o*CB NP@EXOs have the potential as a biomimetic PTT agent, which has remarkable anticancer activity by promoting the apoptosis of tumor cells in mice.

**Figure 6.**
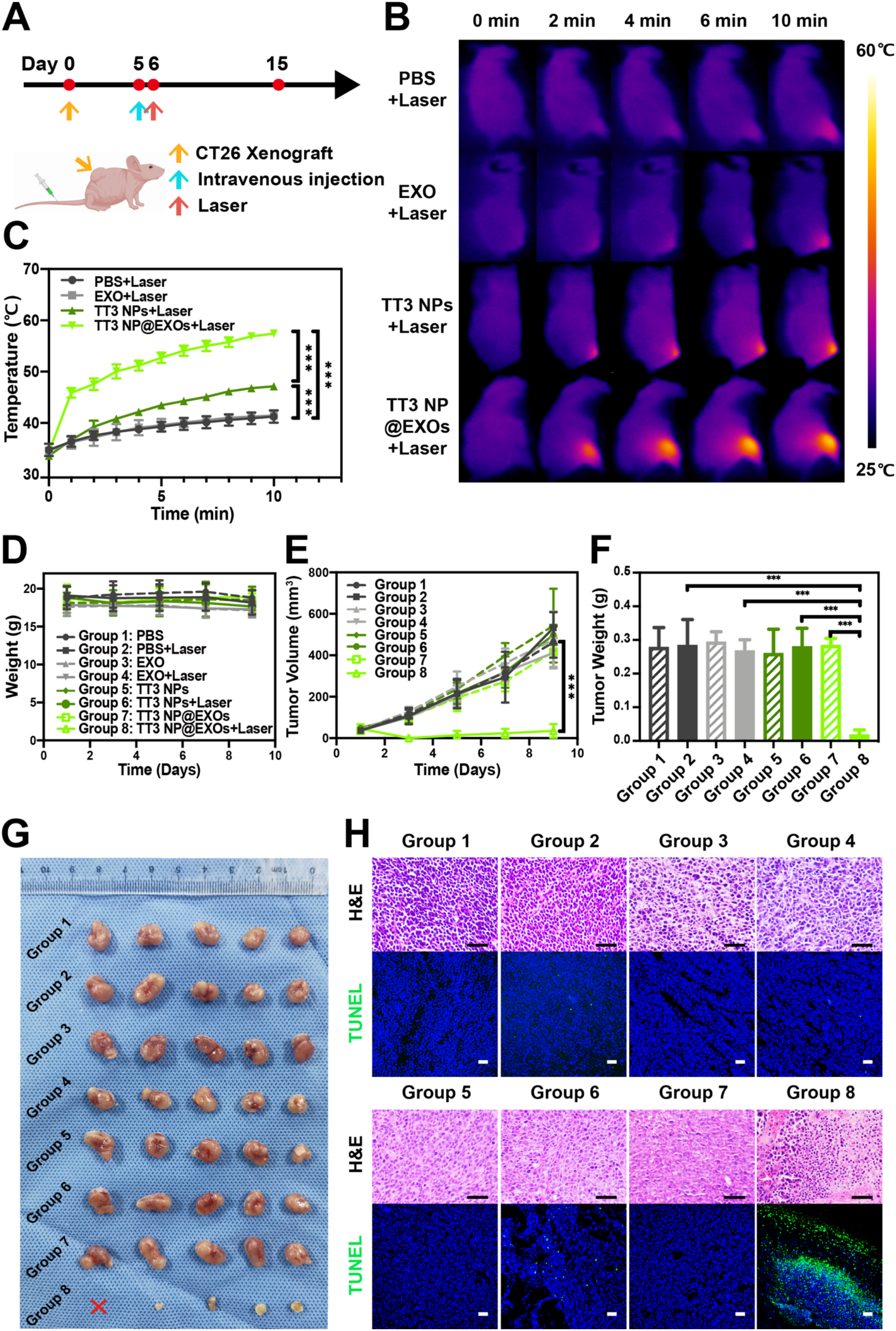
*In vivo* PTT on CT26 tumor-bearing mice. (A) Flow chart of the PTT on CT26 tumor-bearing mice. CT26 cells were subcutaneously injected on day 0, the materials (1 mg mL^-1^, 200 μL) were intravenously injected on day 5 when the tumor volumes had reached approximately 40–60 mm^3^, and PTT was performed on day 6. On day 15, all the mice were sacrificed for further analysis. (B) Infrared thermal images of mice intravenously injected with PBS, EXO, TT3-*o*CB NPs and TT3-*o*CB NP@EXOs (1 mg mL^-1^, 200 μL, termed TT3 NPs and TT3 NP@EXOs, respectively), followed by laser irradiation (808 nm, 1.0 W cm^-2^) 24 h post injection for 0 min, 2 min, 4 min, 6 min and 10 min. (C) Temperature curves of the subcutaneous tumor sites of the mice intravenously injected with EXO, PBS, TT3-*o*CB NPs and TT3-*o*CB NP@EXOs (1 mg mL^-1^, 200 μL) with continuous laser irradiation (808 nm, 1.0 W cm^-2^, 10 min). Body weights (D), relative tumor volumes (E) and tumor weights (F) of the mice after different treatments (n=5, \*\*\**P*< 0.001). (G) Images of tumors collected on day 15 from mice in different groups. (H) H&E and TUNEL staining of tumor tissues from mice in different groups, scale bar: 50 μm (group 1: PBS, group 2: PBS+laser, group 3: EXO, group 4: EXO+laser, group 5: TT3-*o*CB NPs, group 6: TT3-*o*CB NPs+laser, group 7: TT3-*o*CB NP@EXOs, group 8: TT3-*o*CB NP@EXOs+laser).

### 2.7 Toxicity and biosafety of TT3-*o*CB NP@EXOs in CT26 tumor-bearing mice

In addition, toxicity and biocompatibility are critical considerations for AIE NPs used *in vivo*. So, mice were sacrificed one month post the injection of TT3-*o*CB NP@EXOs, and their vital organs were further evaluated via histological and blood biochemistry analyses. We observed no visible organ damage or inflammatory lesions in any of the vital organs (heart, liver, spleen, lung, kidney and gut, Figure S10, Supporting Information). Moreover, no measurable hepatic or renal damage was observed (Figure S11, Supporting Information). To investigate the *in vivo* fate of the TT3-*o*CB NP@EXOs, we performed NIR-II fluorescence imaging of the *ex vivo* organs for the detection of residual NPs, and found that only the livers and spleens of the TT3-*o*CB NP@EXO treated-mice exhibited a weak fluorescence signal 30 days post the injection (Figure S12, Supporting Information). Our previous study indicated that the long aliphatic chains of the AIE molecule significantly enhance the hepatobiliary excretion of the molecule encapsulated micelles^[28]^. Owing to the long aliphatic chains possessed by TT3-*o*CB NPs (Figure S1, Supporting Information), TT3-*o*CB NP@EXOs may be eliminated by the hepatobiliary excretion. In summary, our results illustrated that TT3-*o*CB NP@EXOs had very good biocompatibility and potential for biomedical theranostic applications.

## 3. Conclusions

In this study, we successfully developed a kind of hybrid nanovesicles by combining tumor cell-derived EXO and NIR-II fluorescent organic AIE NPs to specifically kill tumor cells via PTT treatment. Compared with TT3-*o*CB NPs, TT3-*o*CB NP@EXOs displayed excellent biocompatibility and prolonged blood circulation. Thus, the TT3-*o*CB NP@EXOs could specifically target homologous tumors *in vivo*, based on the homing effect arising from the similar membrane-specific protein profiles between tumor cells and their derived-EXO. Because of their high NIR-II fluorescence brightness and excellent photostability, TT3-*o*CB NP@EXOs were excellent NIR-II fluorophores and have great potential for use in diagnostics. Additionally, TT3-*o*CB NP@EXOs had the potential for use as biomimetic NPs for NIR-II imaging-guided PTT due to their high and stable photothermal conversion capacity under the irradiation of 808 nm laser. Therefore, we firstly developed the tumor cell-derived EXO/organic AIE NP hybrid nanovesicles for NIR-II fluorescence imaging guided-PTT. This study provides a universal strategy for increasing the specific tumor targeting capability of AIE NPs. The EXO mediated NP delivery platform is attractive and the future research may focus on improving this delivery platform to achieve a better treatment efficacy. The biomimetic AIE NPs can be utilized for combinatory strategy of diagnosis and PTT in clinical trials in the near future.

## 4. Experimental Section

### Materials

Pluronic F-127 (P2443) was bought from Sigma-Aldrich, USA. Dulbecco’s modified eagle medium (DMEM, C11995500BT) and fetal bovine serum (FBS, 10099141) were bought from Gibco, USA. Phosphate buffered saline (PBS, P1020) and the hematoxylin and eosin staining kit (G1120) were purchased from Solarbio, China. Hoechst 33342 reagent (100 ×, C1022) was purchased from Beyotime, China. The BCA kit (MA0082) and 4% neutral buffered formalin (MA0192) were bought from Meilunbio, China. All the materials above were used directly without further purification.

### Cell lines and cell culture

The murine colorectal cancer cell line CT26, murine liver cancer cell line Hep1-6 and murine fibroblast cell line NIH3T3 were purchased from the cell bank of the Chinese Academy of Sciences (Shanghai, China). The CT26, Hep1-6 and NIH3T3 cells were cultured in DMEM containing 10% FBS in a 5% CO_2_ incubator at 37 °C (Thermo, USA).

### Isolation of tumor cell-derived EXO and assembly of TT3-oCB NP@EXOs

CT26 cells were grown in T75 flasks and cultured in DMEM until 100% confluence. To isolate EXO, the culture medium was collected and centrifuged at 200 × g for 10 min, 2000 × g for 10 min and 10000 × g for 30 min separately to remove cells and debris. Then, CT26 EXO were isolated by ultracentrifugation at 100000 × g for 60 min at 4 °C. Electroporation was then carried out to transfer TT3-*o*CB NPs into EXO at 250 V and 350 μF on a Gemini X2 electroporation system (BTX, USA). After electroporation, the mixture was incubated at 37 °C for 30 min to allow the recovery of the electroporated EXO membrane. Then, unpackaged TT3-*o*CB NPs were removed by ultracentrifugation (100000 × g, 60 min, 4 °C), and the precipitate was retained and dissolved in PBS, thereby yielding TT3-*o*CB NP@EXOs for further use.

### Characterizations of TT3-oCB NP@EXOs

TEM images were acquired by a Tecnai G2 Spirit TEM microscope at 120 kV (FEI, USA). The DLS and zeta potential values were measured on a Malvern Zetasizer Nano instrument (Malvern, UK), and the absorption spectra were recorded by a UV-2000 photospectrometer with UVProbe v2.42 (Shimadzu, Japan). NIR-II PL spectra were collected by a FLS980 spectrometer (Edinburgh Instruments, UK). To measure the photothermal effects of the solutions (0.5 mg ml^-1^, 100 μL), images were acquired by a TiS20 infrared camera (Fluke, USA), and an 808 nm laser was then applied at different power densities. To assess photothermal stability, five irradiation-cooling cycles were implemented with the 808 nm laser (1.0 W cm^-2^).

### Bright field and fluorescence microscopic imaging

Both the bright field and fluorescence microscopic imaging systems were based on a microscope (NIR II-MS, Sunny Optical, China) equipped with an oil-immersion objective (60×, Nikon CFI Plan Fluor) and a silicon-based camera (GA1280, TEKWIN SYSTEM, China). For bright field imaging, the white light used as the light source was transilluminated on the sample, and was collected by the objective and recorded by the camera. For visible fluorescence microscopic imaging, a mercury lamp was used as the excitation source. Passing through a 375/28 nm bandpass filter, reflected by a 415 nm long-pass (LP) dichroic mirror and passing through the objective, the excitation beam was illuminated onto the sample. The excited visible fluorescence from Hoechst dye was collected by the same objective, passed through the same 415 nm LP dichroic mirror and a 460/60 nm bandpass filter, and finally recorded by the aforementioned camera. For NIR-II fluorescence microscopic imaging, an 808 nm laser was used as the excitation source. Reflected by a 900 nm LP dichroic mirror (DMLP900R, Thorlabs, USA) and passing through the aforementioned objective, the 808 nm laser illuminated onto the sample. The excited NIR-II fluorescence was collected by the objective, passed through the same 900 nm LP dichroic mirror and a 900 nm long-pass filter (FELH0900, Thorlabs, USA), and finally recorded by the camera. For cell sample preparation, CT26, Hep1-6 and NIH3T3 cells were smeared onto glass slides in 24-well plates (2000 cells per well) and incubated with PBS, EXO (50 μg mL^-1^, measured by BCA assays), TT3-*o*CB NPs (50 μg mL^-1^) or TT3-*o*CB NP@EXOs (50 μg mL^-1^ for TT3-*o*CB NPs) for 24 h. Then, the slides were washed with PBS three times and counterstained with Hoechst 33342 reagent (100 ×, C1022, Beyotime, China) for 5 min. After washing with PBS three times, the slides were observed by the bright field and fluorescence microscope.

### Optical system for in vivo NIR-II fluorescence imaging

The biodistributions of TT3-*o*CB NPs and TT3-*o*CB NP@EXOs *in vivo* were determined with a lab-built NIR-II fluorescence imaging system equipped with an 808 nm laser. The laser beam was coupled with an optical collimator and expended by a lens to provide uniform illumination in the irradiated area. An InGaAs camera equipped with a prime lens (focal length: 35 mm, antireflection coating: 900-1700 nm, TEKWIN SYSTEM, China) was utilized to detect the fluorescence signals with 1100 nm long-pass filters (FELH1100, Thorlabs, USA).

### In vitro PTT study

CT26, Hep1-6 and NIH3T3 cells were seeded in 96-well plates (10000 cells per well with 100 μL suspension) and cultured at 37 °C in a 5% CO_2_ incubator for 12 h. EXO (50 μg mL^-1^, measured by BCA assays), TT3-*o*CB NPs (50 μg mL^-1^) and TT3-*o*CB NP@EXOs (50 μg mL^-1^ for TT3-*o*CB NPs) were added to the cells and incubated for 24 h, followed by treatment with or without 808 nm laser irradiation (1.0 W cm^-2^, 5 min). The cell viability was then determined by the MTS assay.

### In vivo NIR-II fluorescence imaging

A total of 200 μg TT3-*o*CB NPs or TT3-*o*CB NP@EXOs resuspended in 200 μL of PBS was intravenously administered into different groups of mice (three CT26 tumor-bearing mice per group). Then, the mice in different groups were anesthetized and imaged using the aforementioned *in vivo* NIR-II fluorescence imaging system. For the time-dependent biodistribution study, blood samples, main organs and tumors were harvested from the mice, which were sacrificed at selected time points after administration (12 h, 24 h, 48 h and 72 h), and imaged using the same NIR-II fluorescence imaging system. The *in vivo* study was approved by the Animal Use and Care Committee at Zhejiang University.

### In vivo PTT study

When the tumor volumes reached approximately 40-60 mm^3^, the mice were ready for PTT and randomly divided into eight groups (five mice per group): (1) PBS only; (2) PBS + 808 nm laser (1 W cm^-2^, 10 min); (3) EXO (1 mg mL^-1^, 200 μL) only; (4) EXO (1 mg mL^-1^, 200 μL) + 808 nm laser; (5) TT3-*o*CB NPs (1 mg mL^-1^, 200 μL) only; (6) TT3-*o*CB NPs (1 mg mL^-1^, 200 μL) + 808 nm laser; (7) TT3-*o*CB NP@EXOs (1 mg mL^-1^ for TT3-*o*CB NPs, 200 μL) only; and (8) TT3-*o*CB NP@EXOs (1 mg mL^-1^ for TT3-*o*CB NPs, 200 μL) + 808 nm laser. The mice in the laser groups were anesthetized and then irradiated with an 808 nm laser (1.0 W cm^2^, 10 min) 24 h post injection of the dispersions in different groups. Tumor growth was assessed by measuring the xenograft sizes, and the tumor volumes were calculated every two days with the following formula: volume = (length × width^2^)/2. The tumor and body weights were observed and recorded at the same time intervals. The infrared thermal imaging and temperature increases were monitored by a TiS20 infrared camera (Fluke, USA). These mice were monitored until sacrifice on day 15.

### Histological analysis

Tumors from mice in all groups were harvested after 24 h. They were fixed with 4% neutral buffered formalin, embedded in paraffin, sliced, as well as stained with hematoxylin and eosin, according to the manufacturer’s manual, and observed under an inverted optical microscope (Zeiss, Germany).

### Statistical analysis

ImageJ 32 software was used to quantify the fluorescence signals in the *in vivo* biodistribution study, and values were normalized to the maximum value at 24 h for each group. The quantitative data were analyzed by Student’s *t*-test. *P* values denote differences between the control and experimental samples, and \*\*\**P* < 0.001, \*\**P* < 0.01 and \**P* < 0.05 indicate statistical significance.

## Supporting Information

Supporting Information is available from the Wiley Online Library or from the author.

## Acknowledgements

Y.L. (Yirun L.), X.F. and Y.L. (Yuanyuan L.) contributed equally to this work. This work was supported by National Key Research and Development Project (2017YFC0110802), Zhejiang province Key Research and Development Project (2020C01059), National Natural Science Foundation of China (81872297, 81874059, 61975172 and 61735016), Zhejiang Engineering Research Center of Cognitive Healthcare (2017E10011), National Key Scientific Instrument and Equipment Development Project (81827804), and Fundamental Research Funds for the Central Universities (2020-KYY-511108-0007).

## Data Availability Statement

Research data are not shared.

---

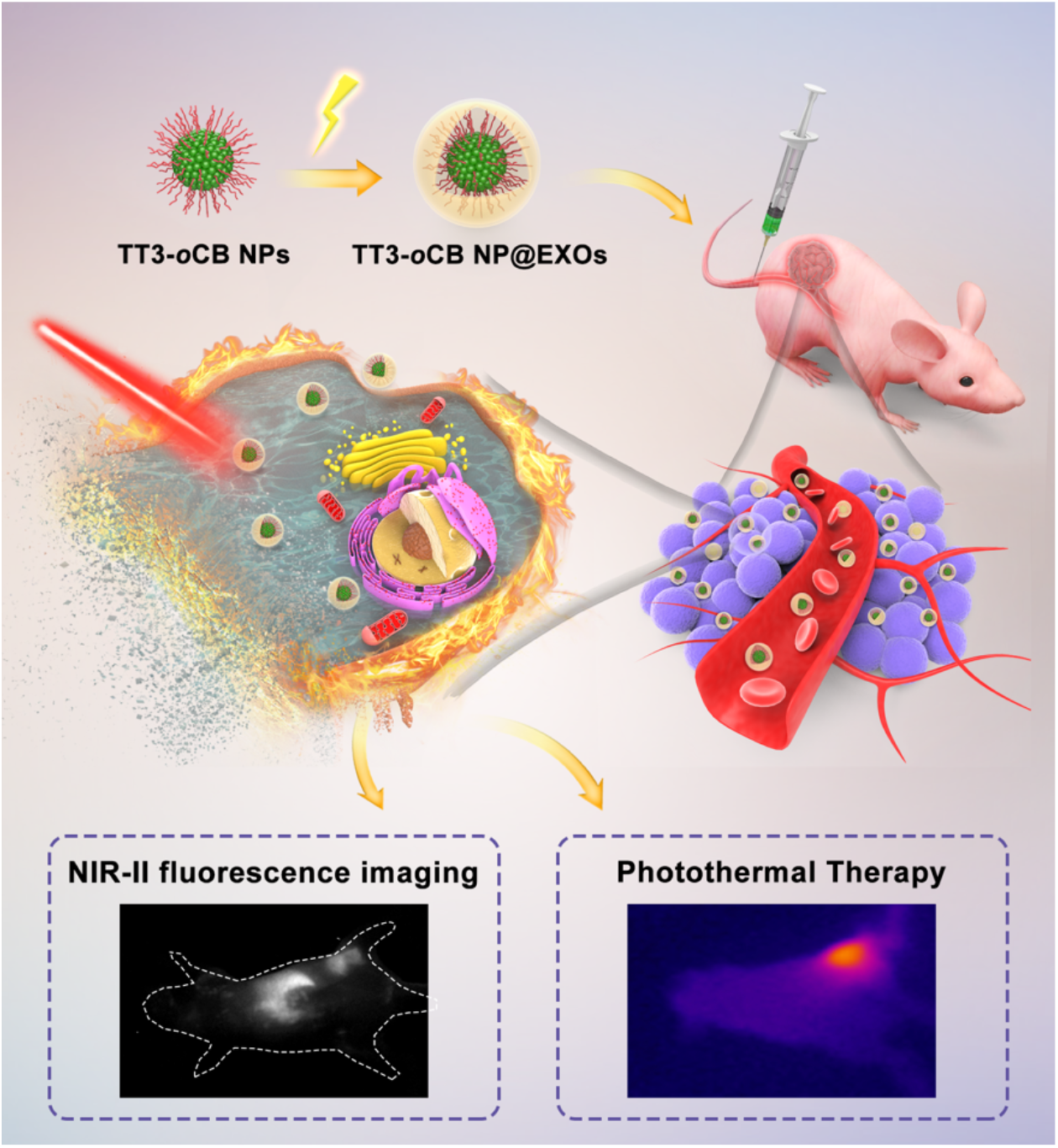

