## Supporting Information for "AIE nanoparticles camouflaged with tumor cell-derived exosomes for NIR-II imaging-guided photothermal therapy"

Prof. Hui Lin

Zhejiang Engineering Research Center of Cognitive Healthcare, Sir Run Run Shaw Hospital, School of Medicine, Zhejiang University, 310016, China.

Dr. Xiaoxiao Fan, Runze Chen, Huwei Ni, Zhe Feng, Prof. Jun Qian

State Key Laboratory of Modern Optical Instrumentations, Centre for Optical and Electromagnetic Research, College of Optical Science and Engineering, International

Research Center for Advanced Photonics, Zhejiang University, Hangzhou 310058, China.

Dr. Yuanyuan Li

College of Veterinary Medicine, Jilin University, Changchun 130062, China

Prof. Ben Zhong Tang

Shenzhen Institute of Aggregate Science and Technology, School of Science and Engineering, The Chinese University of Hong Kong, Shenzhen, Guangdong 518172, China

### Methods

**Western blot experiments**: Cells were seeded in 6 cm dishes and lysed with lysis buffer, and the supernatant was harvested after centrifugation. Then, protein quantitation was performed by the BCA method (MA0082, Meilunbio, China). After adding SDS loading buffer (MA0003, Meilunbio, China), the samples were boiled for 5 min, separated by 10% SDS-PAGE and transferred to a PVDF membrane (IPVH00010, Millipore, USA). The PVDF membrane was blocked with blocking buffer for 60 min, washed with TBST (CW0043, CWbio, China) three times, incubated with a primary antibody (anti-TSG101 ab125011, anti-CD9 ab92726, anti-CD81 ab109201, anti-beta actin, anti-Bax ab182733, anti-cleaved caspase 3 ab32042, anti-

Bcl2 ab32124 and anti-tubulin ab6046, Abcam, USA) at 4 °C overnight, and further incubated with a secondary antibody for 1 h. Finally, the protein bands were detected by a ChemiDoc Touch Imaging System (Bio-Rad, USA).

**In vitro cytotoxicity assays:** CT26, Hep1-6 and NIH3T3 cells were incubated in 96-well plates (5000 cells per well with 100 μL of suspension) for 12 h. Then, the culture medium was replaced with 100 μL fresh culture medium containing TT3-*o*CB NPs or TT3-*o*CB NP@EXOs at different concentrations (0, 2, 5, 10, 25, 50 μg $mL^{-1}$). The cells were incubated for an additional 24 h, and cell viability was then assessed by the MTS assay (G3580, Promega, USA). After the culture medium was removed, the wells were washed three times with PBS, and fresh medium containing MTS reagent (10%) was added to each well. The cells were then incubated for another 4 h, and the absorbance of the culture medium at 490 nm was measured by a multidetection microplate reader (Thermo, USA) to evaluate the cell viability.

**Clone formation assays:** CT26, Hep1-6 and NIH3T3 cells were seeded in 6-well plates at a concentration of 5000 cells $well^{-1}$ and cultured at 37 °C in a 5% $CO_2$ incubator for 12 h. EXO (50 μg $mL^{-1}$, measured by BCA assays), TT3-*o*CB NPs (50 μg $mL^{-1}$) and TT3-*o*CB NP@EXOs (50 μg $mL^{-1}$ for TT3-*o*CB NPs) were added to the cells and incubated for 24 h, followed by the 808 nm laser irradiation (1.0 W $cm^{-2}$, 5 min). After 14 days of culture, visible colonies fixed in 4% paraformaldehyde were stained with crystal violet solution (C0121, Beyotime, China) and photographed.

**TUNEL staining analysis:** TUNEL staining was performed using a TUNEL BrightGreen Apoptosis Detection Kit to identify apoptotic cells (A112-01, Vazyme, China). Samples were prepared according to the manufacturer's manual. Briefly, CT26, Hep1-6 and NIH3T3 cells were smeared onto glass slides (5000 cells per well in a 24-well plate), incubated with NPs (50 μg $mL^{-1}$) for 24 h, and subjected to the 808 nm laser irradiation (1.0 W $cm^{-2}$, 5 min). The cells were washed with PBS twice, fixed with 4% paraformaldehyde at 4 °C for 25 min, and then treated with 0.2% Triton® X-100 solution for 5 min. After equilibration for 30 min, the slides were incubated with TdT solution for 1 h at 37 °C and counterstained with Hoechst 33342 reagent for 5 min. TUNEL-positive cells were imaged with a fluorescence microscope (ZEISS, Germany), and cells with green fluorescence were considered to be apoptotic.

Paraffin-embedded tumor sections of CT26 tumor-bearing mice from different groups were deparaffinized with xylene, rehydrated, and permeabilized with proteinase K solution (20 μg $mL^{-1}$, ST532, Beyotime, China) for 20 min at room temperature. The next procedures were performed as described above for the TUNEL staining of cells.

**Flow cytometry study *in vitro*:** Apoptosis was measured by flow cytometry using an Annexin V-FITC/propidium iodide (PI) apoptosis kit (70-AT101-60, MultiSciences, China). CT26, Hep1-6 and NIH3T3 cells were plated on 12-well plates (50000 cells per well). After incubation with NPs (50 μg $mL^{-1}$) for 24 h, the cells were irradiated with an 808 nm laser (1.0 W $cm^{-2}$, 5 min), harvested and resuspended in 500 μL 1 ×

binding buffer. Then, 5 μL Annexin V-FITC and 10 μL PI were added to each well containing the cell suspension and incubated for 5 min at room temperature in the dark. The specimens labeled with Annexin V-FITC and PI were detected by a FACSCanto II flow cytometer (BD Biosciences, USA), and the data were analyzed using Flow Jo V10.

**Wound healing assays:** Wound healing assays were used to evaluate the effects of PBS, EXO, TT3-*o*CB NPs and TT3-*o*CB NP@EXOs on the migration of CT26, Hep1-6 and NIH3T3 cells. CT26, Hep1-6 and NIH3T3 cells were incubated with PBS, EXO, TT3-*o*CB NPs and TT3-*o*CB NP@EXOs (same concentration: 1 mg $mL^{-1}$) for 24 h. Then, the cells were wounded by the dragging of a 200 μL pipet tip, and bright field images were acquired immediately. After irradiation (808 nm, 1 W $cm^{-2}$, 5 min), the status of wound closure was recorded by inverted microscopy 48 h post irradiation.

**Measurement of the photothermal conversion efficiency:** The TT3-*o*CB NP and TT3-*o*CB NP@EXO (100 μL, 0.1 mg $ml^{-1}$) aqueous dispersions were irradiated by an 808 nm laser (0.5 W $cm^{-2}$) for a prolonged period until the temperature reached a plateau, and then cooled naturally. The changes in temperature were recorded every ten seconds until cooling to room temperature was achieved. The photothermal conversion efficiency ($\eta$) was estimated according to the following equation[1]:

$$\eta = \frac{hS(T_{max} - T_{surr}) - Q_{dis}}{I(1 - 10^{-A_{808}})}$$

where $T_{max}$ represents the maximum temperature, $T_{surr}$ represents the room temperature, $Q_{dis}$ represents the heat dissipation caused by the light absorption of the container, $I$ represents the power of the laser (0.3 W), and $A_{808}$ represents the absorbance of NPs at 808 nm.

The $hS$ value ($h$ is the heat transfer coefficient and $S$ is the container's surface area) can be calculated by the following equation:

$$hS = \frac{m_D C_D}{\tau_S}$$

where $m_D$ is the mass of aqueously dispersed NPs (0.1 g), $C_D$ is the heat capacity of the solvent (water, 4.2 J $g^{-1}$), and $\tau_s$ is the time constant.

**Establishment of xenograft tumors in nude Mice***:* CT26 cells (1000000 cells per mouse) were injected subcutaneously into the right hind legs of female BALB/c nude mice (4 weeks old) to establish a tumor-bearing mouse model. The study was approved by the Animal Use and Care Committee at Zhejiang University.

***In vivo* biodistribution and biosafety of TT3-*o*CB NP@EXOs:** A total of 200 μg TT3-*o*CB NP@EXOs resuspended in 200 μL of PBS was intravenously administered to mice in different groups (three ICR mice per group). For the control group, ICR mice were intravenously injected with PBS (200 μL, 1×). After one month, all the mice were sacrificed. Blood samples were collected by cardiac puncture, and hematology studies were performed. The vital organs (heart, liver, spleen, lung, kidney and gut) of the mice were surgically dissected and subjected to NIR-II fluorescence imaging, prior to being fixed with 4% paraformaldehyde for 24 h, embedded in paraffin, sliced, as well as stained with hematoxylin and eosin according to the manufacturer's manual, and observed under an inverted optical microscope (Zeiss, Germany).

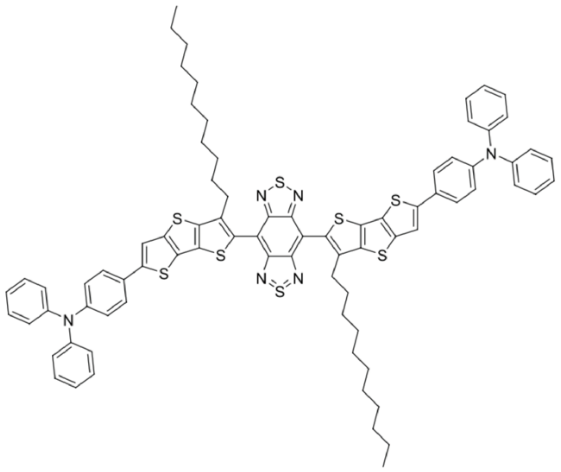


**Figure S1.** Molecular structure of the TT3-*o*CB compound.

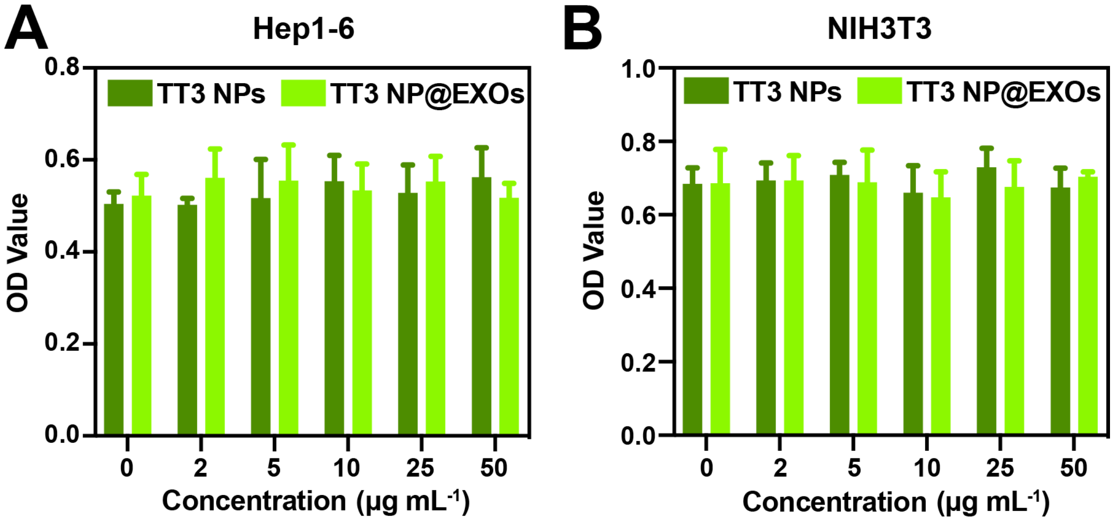


**Figure S2.** *In vitro* viability of Hep1-6 and NIH3T3 cells incubated with TT3-*o*CB NPs and TT3-*o*CB NP@EXOs (termed TT3 NPs and TT3 NP@EXOs, respectively) at different concentrations for 24 h.

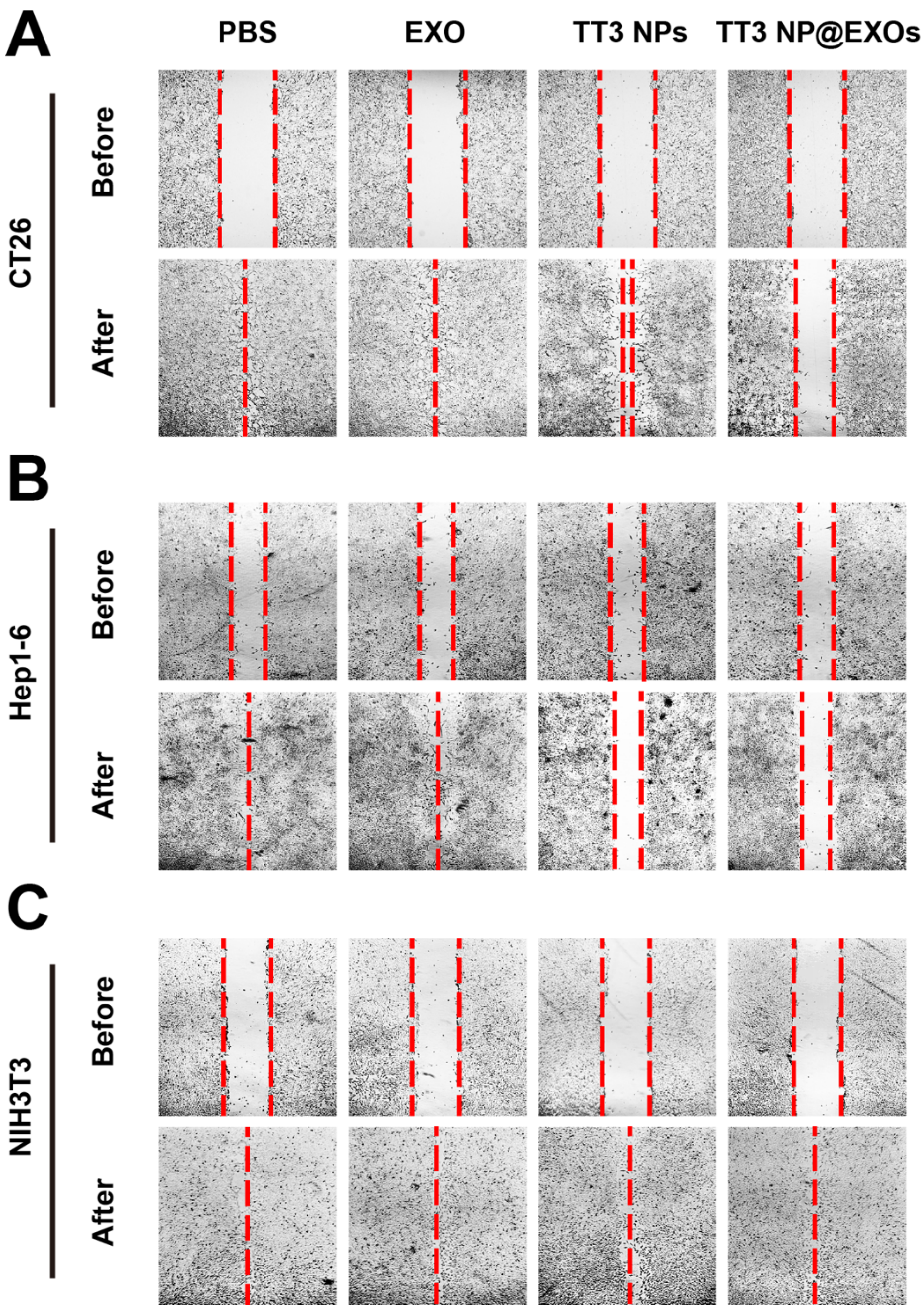


**Figure S3.** Wound healing assays of CT26, Hep1-6 and NIH3T3 cells incubated with PBS, EXO, TT3-*o*CB NPs and TT3-*o*CB NP@EXOs (termed TT3 NPs and TT3 NP@EXOs, respectively) for 24 h and irradiated by an 808 nm laser (1.0 W $cm^{-2}$, 5 min).

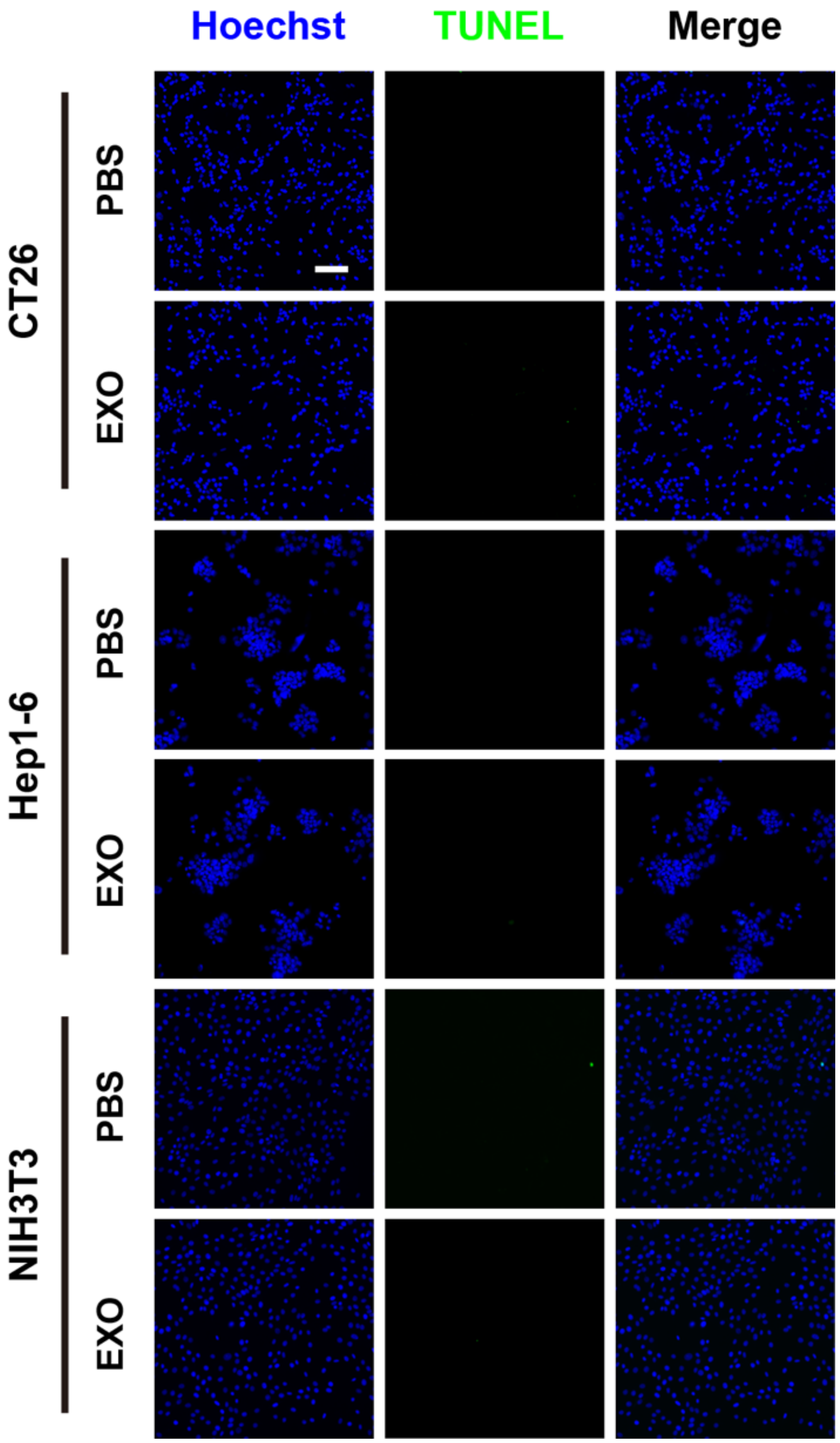


**Figure S4.** TUNEL staining of CT26, Hep1-6 and NIH3T3 cells incubated with PBS and EXO (1 mg $mL^{-1}$), under the irradiation of an 808 nm laser (1.0 W $cm^{-2}$, 5 min; green: TUNEL-positive cells, blue: Hoechst-stained cell nuclei). Scale bar: 50 μm.

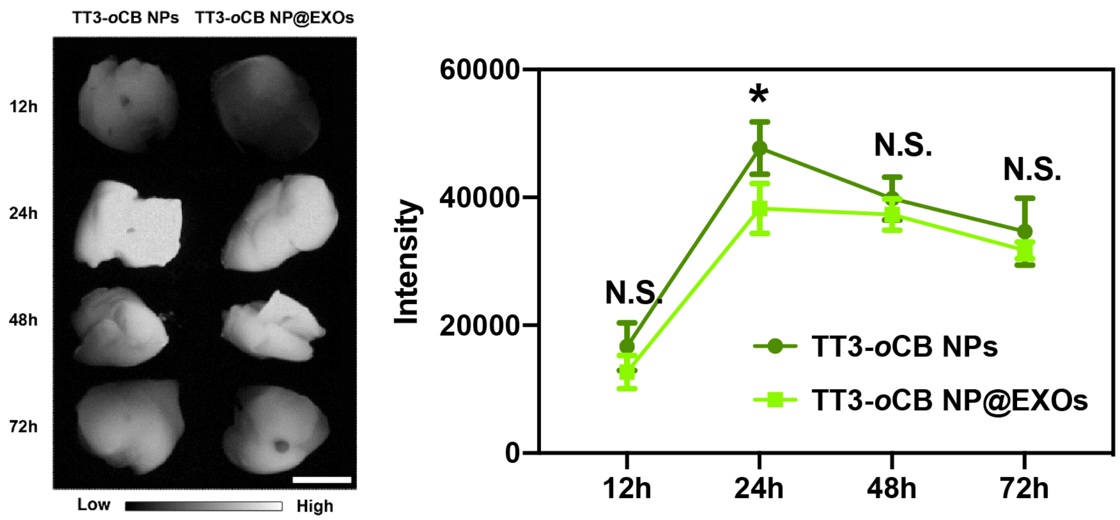


**Figure S5.** Time-dependent NIR-II fluorescence images (left) and intensity changes (right) of the livers in the TT3-*o*CB NP- and TT3-*o*CB NP@EXO-treated groups, 12, 24, 48, and 72 h post injection (n = 3, N.S. $P > 0.05$, *$P < 0.05$, scale bar: 10 mm).

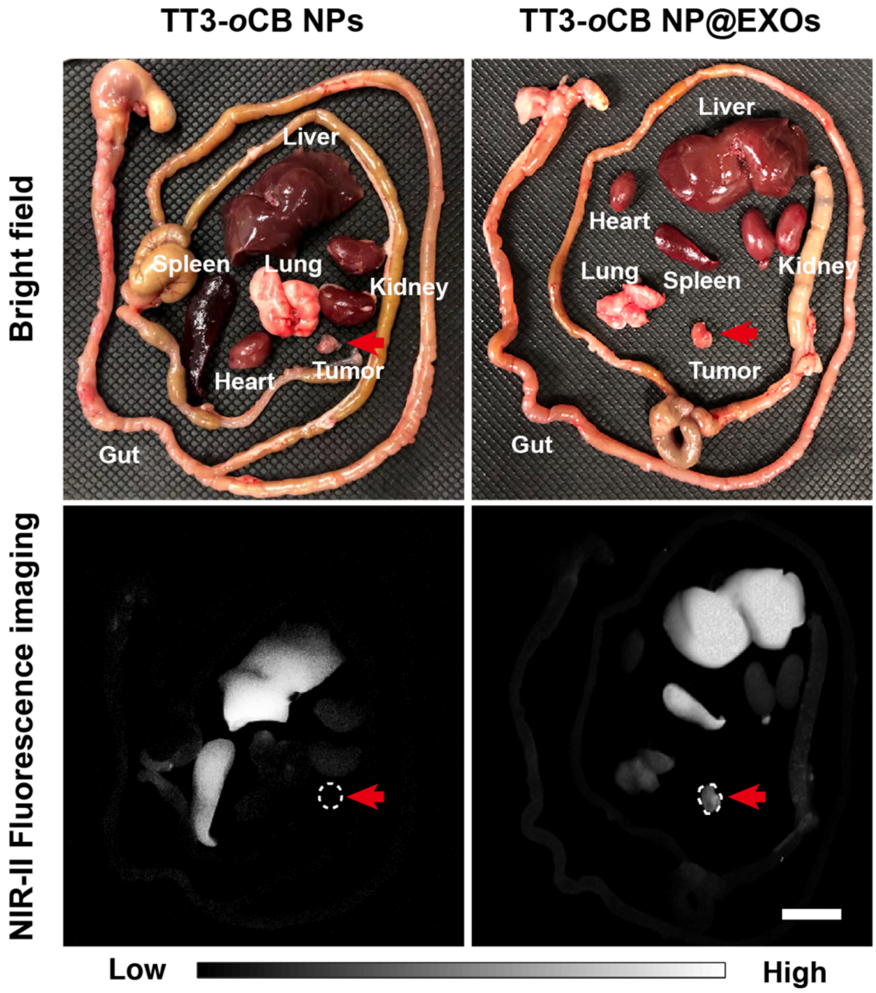

**Figure S6.** *Ex vivo* bright field and NIR-II fluorescence images of tumors and vital organs from CT26 tumor-bearing mice 24 h after the intravenous (i.v.) administration of TT3-*o*CB NPs (1 mg mL$^{-1}$, 200 μL, left) and TT3-*o*CB NP@EXOs (1 mg mL$^{-1}$, 200 μL, right) (the red arrows indicate tumors). Scale bar: 10 mm.

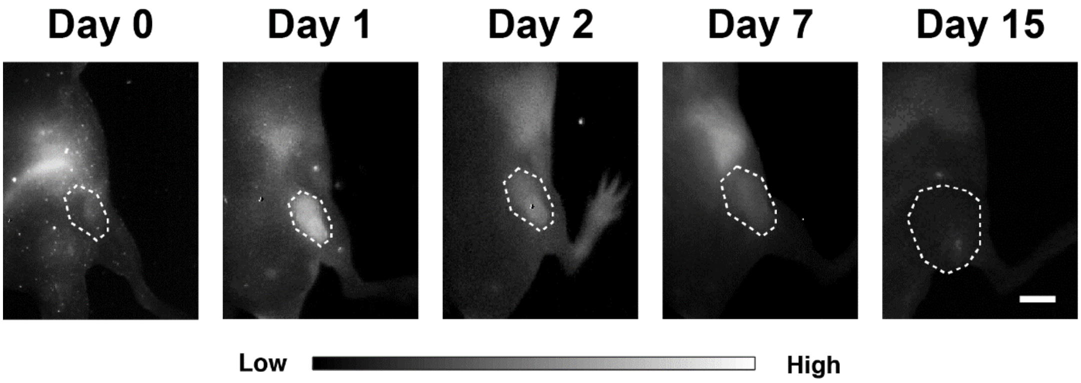


**Figure S7.** *In vivo* long-term tracing of the subcutaneous CT26 tumor on nude mouse treated with TT3-*o*CB NP@EXOs, using NIR-II fluorescence imaging. Scale bar: 10 mm.

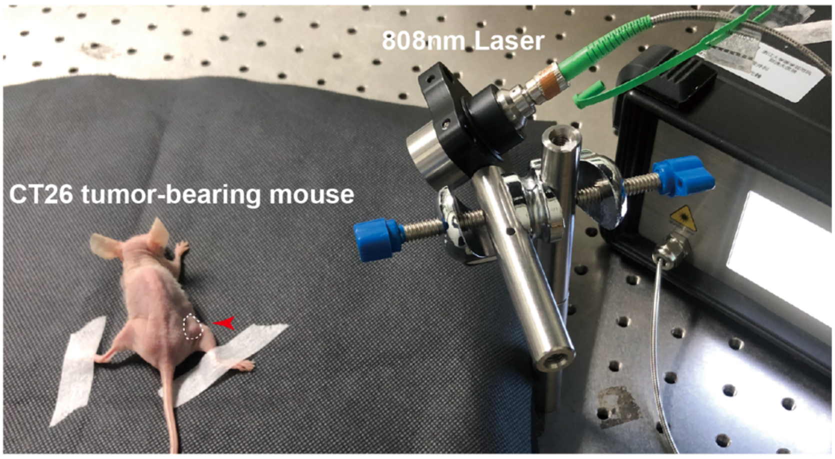


**Figure S8.** Schematic illustration of the PTT system for irradiating CT26 tumor-bearing mice with an 808 nm laser (1.0 W cm$^{-2}$, 10 min).

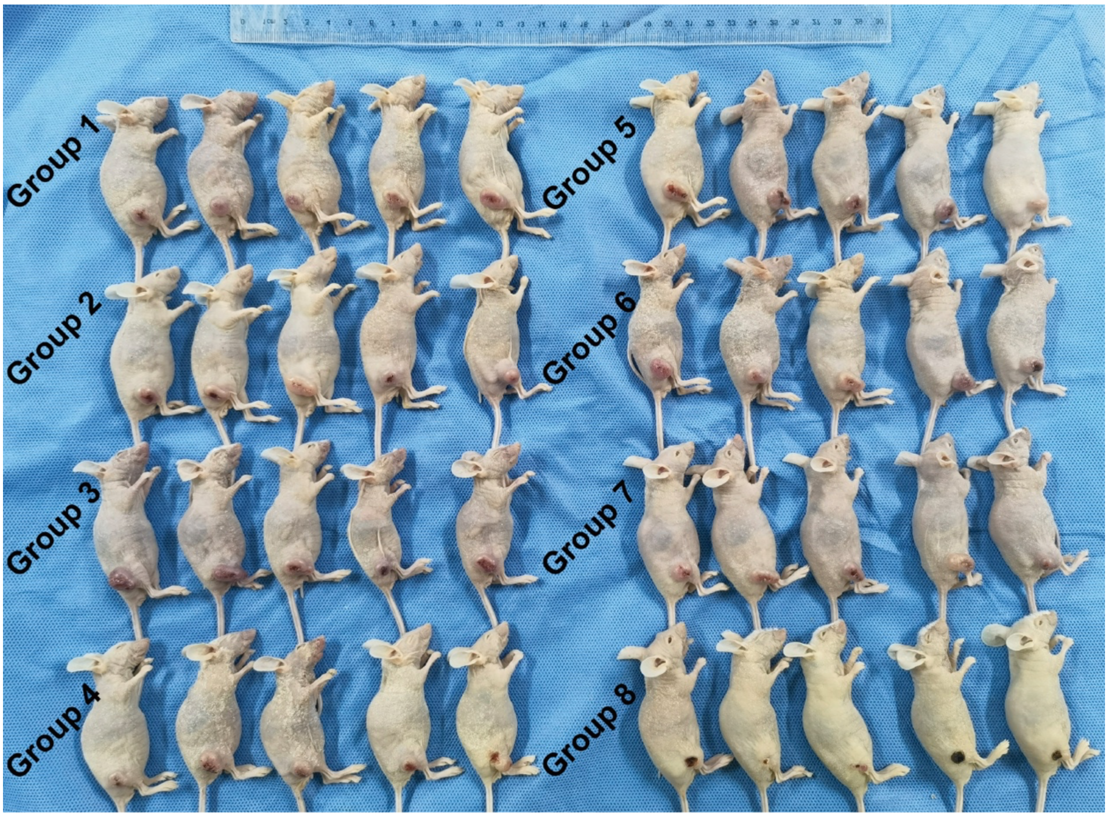


**Figure S9.** Images of CT26 tumor-bearing mice from different groups on day 15 (five mice per group; group 1: PBS, group 2: PBS+laser, group 3: EXO, group 4: EXO+laser, group 5: TT3-*o*CB NPs, group 6: TT3-*o*CB NPs+laser, group 7: TT3-*o*CB NP@EXOs, group 8: TT3-*o*CB NP@EXOs+laser).

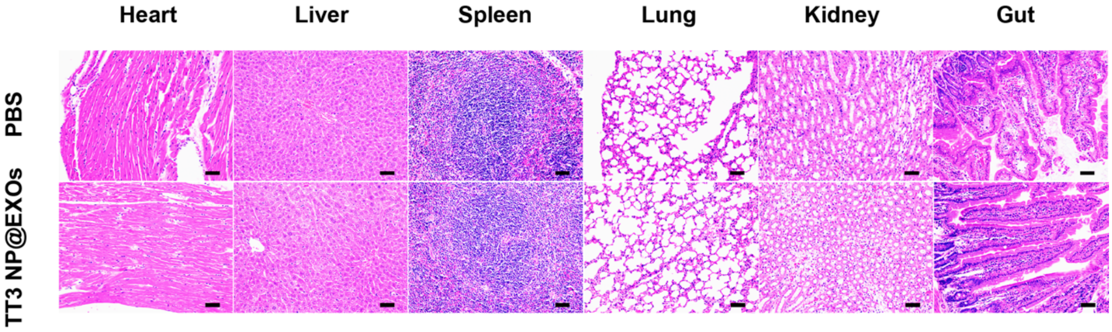


**Figure S10.** H&E staining of vital organs (heart, liver, spleen, lung, kidney and gut) of the mice 1 month after the i.v. administration of PBS (200 μL, upper panel) and TT3-*o*CB NP@EXOs (1 mg $mL^{-1}$, 200 μL, bottom panel). Scale bar: 50 μm.

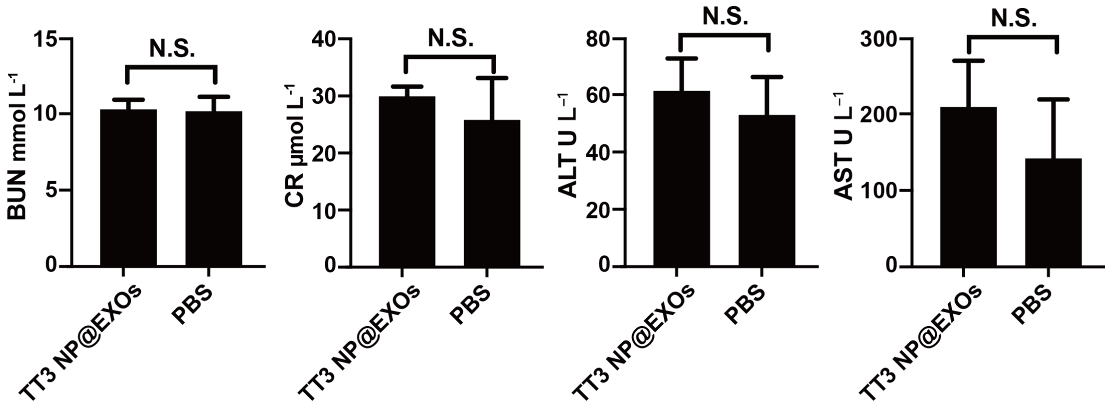


**Figure S11.** Chronic biological toxicity analysis: the hepatic and renal functions of mice treated with and without TT3-*o*CB NP@EXOs for 1 month were assessed (control: PBS, 200 μL, n = 3; experimental group: TT3-*o*CB NP@EXOs: 1 mg mL$^{-1}$, 200 μL, n = 3, N.S. $P > 0.05$).

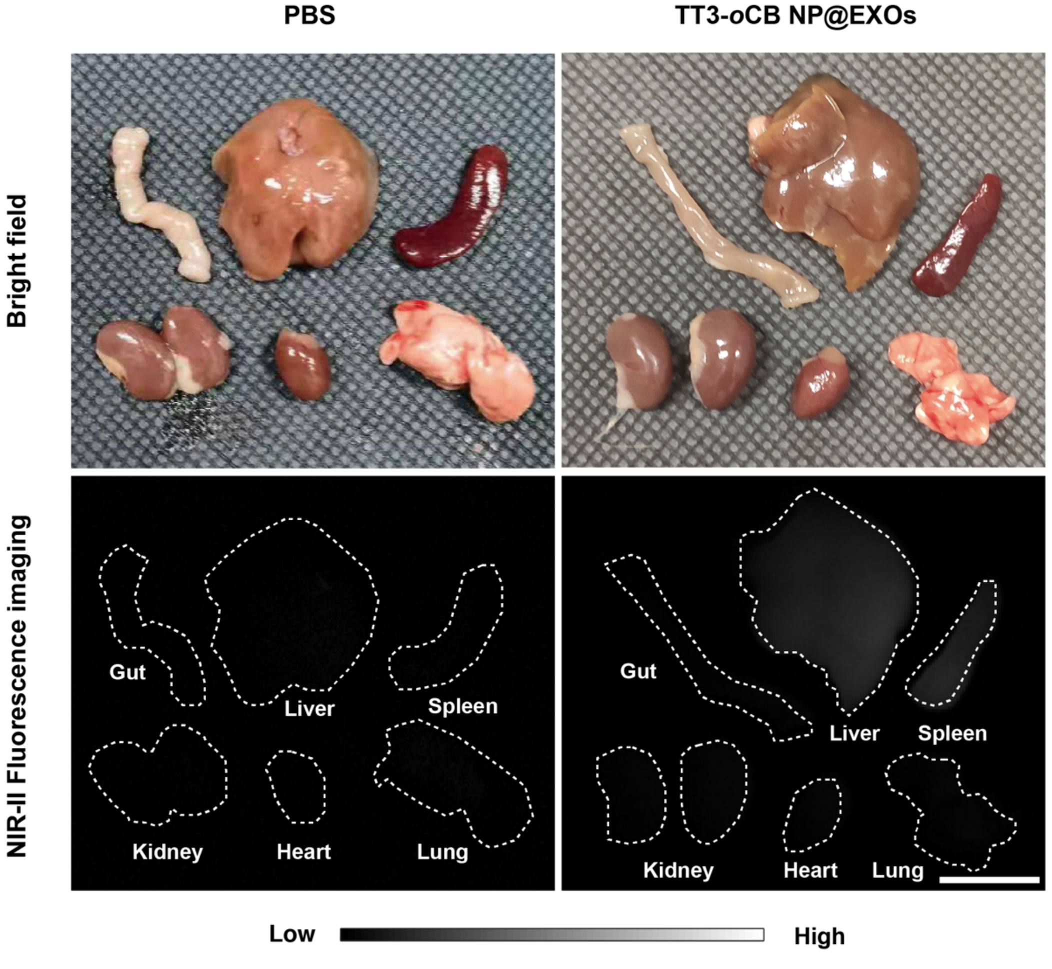


**Figure S12.** Bright field and NIR-II fluorescence images of vital organs from mice 1

month after the i.v. administration of PBS (200 μL, left) and TT3-*o*CB NP@EXOs (1 mg $mL^{-1}$, 200 μL, right). Scale bar: 10 mm.
